# Nitric oxide tunes secreted metabolite bioactivity

**DOI:** 10.1101/2025.11.05.686753

**Authors:** Zachery R. Lonergan, Sarah L. Weisflog, Matthew Scurria, Jinyang Li, Korbinian Thalhammer, Osvaldo Gutierrez, Stuart J. Conway, Dianne K. Newman

## Abstract

The radical nitric oxide (·NO) is short-lived but has imprinted itself on many aspects of physiology and disease. ·NO’s rapid production and consumption, coupled with its intrinsic reactivity, drive its biological importance; thus, defining mechanisms and targets of ·NO reactivity is necessary to assess its fate and impact. Cellular small molecules are a major class of ·NO-reactive targets, possessing a variety of molecular functionalities that can react with ·NO. Yet the capacity for secreted small molecules to react with ·NO, as well as the biological consequences of such reactivity, have received little attention. Here, we explore the reactivity of ·NO with phenazine metabolites, microbially-derived secreted small molecules that possess antibiotic properties and can modulate their microenvironment. Using *Pseudomonas aeruginosa* as a model phenazine producer, we find that ·NO reacts with specific phenazines to yield stable, chemically-distinct products. These chemical transformations significantly attenuate phenazine antibiotic properties, including against the phenazine nonproducer *Staphylococcus aureus*, a competitor with *P. aeruginosa* for niches in the context of infection. By contrast, *P. aeruginosa* experiences rapid loss in viability when phenazines and ·NO react. This toxicity occurs even in the presence of *S. aureus*, which displays resistance to nitrosylated phenazines, implicating a specific toxicity dependent on the formation of the phenazine-NO adduct. These findings highlight the capacity of ·NO to transform metabolite activity and suggest that ·NO can tune microbial interactions in complex environments by a mechanism of action hitherto unappreciated.

## Introduction

Reactive oxygen and nitrogen species are produced continuously in many biological systems. Among these is the radical nitric oxide (·NO), which is widely recognized as a key component of human physiology as a regulator of neurotransmission, vascular dilation, and infection control [1]. ·NO also plays crucial functions for microbial physiology, including as an intermediate during anaerobic respiration and as an inter- and intracellular signaling molecule [2]. The widespread importance of ·NO stems from its intrinsic reactivity coupled with its rapid diffusion, which allows broad-acting effects from a focal point of production over short timescales [1]. ·NO can also react with oxygen and superoxide to yield other reactive nitrogen species, including nitroxyl radicals, nitrogen dioxide, and peroxynitrite, which expand its impact indirectly [3].

While ·NO is commonly described as highly reactive, there is specificity to its chemistry that is intimately linked to its existence as a radical [4]. ·NO preferentially reacts with specific types of molecules and motifs, including phenolic compounds, thiols, amines, and transition metal centers [5–7]. These molecular features are common components of proteins, and significant progress has been made in discerning how ·NO-protein reactivity can alter structure-function relationships [8, 9]. However, these ·NO-reactive motifs are also ubiquitous in the small molecule landscape [10]. Well-described examples demonstrating ·NO-small molecule reactivity include ·NO scavenging via the antioxidant glutathione and ·NO reactivity with the amino acids tyrosine and cysteine [11–13], sugars such as glucose [14], and fatty acids [15]. However, the small molecule landscape is both vast and chemically complex, with ∼10^60^ small molecules theoretically possible [16]. Given their structural diversity and environmental ubiquity, the capacity for ·NO to react with small molecules merits more attention.

Metabolites that are secreted into the environment are common yet often overlooked with respect to their potential to react with ·NO. Like ·NO, secreted small molecules can modify the local chemical environment via their diffusive capacity [17, 18]. An important class of secreted small molecules are phenazine metabolites, whose production has been documented in environments ranging from soils to human chronic infections [19–21]. One prolific phenazine producer is the opportunistic bacterial pathogen *Pseudmonas aeruginosa* that produces the well-studied molecule pyocyanin (PYO), as well as three additional phenazines including 1-hydroxyphenazine (1-OHPhz), phenazine-1-carboxylic acid (PCA), and phenazine-1-carboxyamide (PCN) [22]. Phenazine production plays important roles in *P. aeruginosa* virulence by supporting bacterial anaerobic survival and intoxicating immune cells [23–25], but phenazines also possesses antimicrobial activity, thus serving as natural antibiotics [20, 26, 27]. PYO and other phenazines possess structural features consistent with known ·NO-reactive molecules and are produced in infection environments where ·NO concentrations can be high due to release from innate immune cells [28, 29], but the ability of phenazines to react with ·NO is unclear and disputed [30–32]. We therefore sought to investigate ·NO-phenazine reactivity to clarify the chemistry governing these interactions as an example of how secreted metabolites might impact ·NO fate and biological consequences in disease contexts.

## Results

### Pyocyanin and nitric oxide alter *P. aeruginosa* growth and metabolite detection

·NO is known to inhibit *P. aeruginosa* growth and cause cell death, but the effect of ·NO on physiology is linked to its concentration. Consistent with previous results [33, 34], acute exposure to high concentrations of ·NO kills *P. aeruginosa* (**Fig S1A**), but ·NO also can alter physiology without affecting viability [35]. We found that *P. aeruginosa* can grow with low-level, slow ·NO exposure with no change in the overall cell density (**Fig 1A**). Growth of *P. aeruginosa* also typically results in a blue-green pigmentation of the culture, which largely reflects the production of the secreted phenazine metabolite pyocyanin (PYO) [36]. Regulation of PYO production is complex, but quorum-sensing systems play an important role in facilitating increased production of PYO as cell density increases [37, 38]. Consistent with observations from other strains [39], PYO detection in the presence of subinhibitory levels of NO diminishes over time, without altering growth kinetics or cell density (**Fig 1B**).

**Figure 1.**
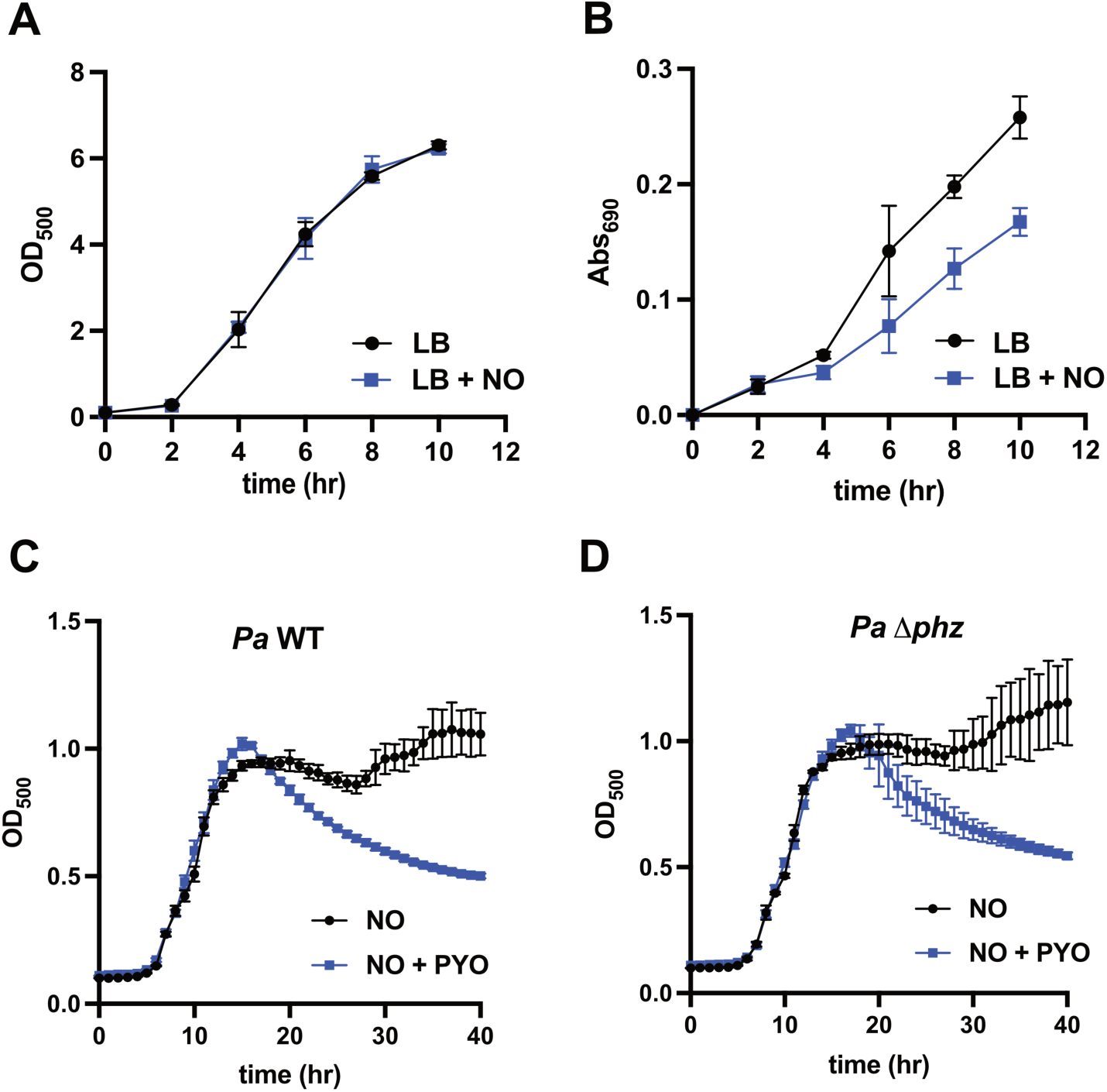
Nitric oxide exposure reduces pyocyanin detection in *P. aeruginosa*. **A)** Growth monitored by absorbance at 500 nm of *P. aeruginosa* (*Pa*) in lysogeny broth (LB) supplemented with 1 mM DETA-NONOate NO donor; n = 3 biological replicates, mean +/− standard deviation (SD). **B)** Relative abundance of pyocyanin (PYO) in supernatant measured via absorbance at 690 nm at times matching (A); n = 3 biological replicates, mean +/− SD. **C)** Representative growth monitored by absorbance at 500 nm of wildtype (WT) *Pa* with 1 mM DETA-NONOate and 100 μM PYO supplemented at the start of growth; n = 3 technical replicates from 1 of 3 biological replicates with mean +/− SD. **D)** Representative growth monitored by absorbance at 500 nm of Δ*phz Pa* with 1 mM DETA-NONOate and 100 μM PYO supplemented at the start of growth; n = 3 technical replicates from 1 of 3 biological replicates, mean +/− SD.

Because PYO detection diminished in the presence of ·NO, we wondered how supplementing growing cultures with excess PYO would further influence *P. aeruginosa* physiology in both a wildtype (WT) strain of *P. aeruginosa*, as well as a phenazine-biosynthesis mutant strain (Δ*phz*) that cannot produce or modify exogenously-provided phenazines [40]. Addition of PYO at physiologically-relevant concentrations does not negatively impact *P. aeruginosa* growth (**Fig S1B,C**). Supplementation of cultures with NO also follows predictable bacterial growth kinetics including a lag phase, exponential growth phase, and stationary phase (**Fig 1C,D**). The combination of PYO and ·NO permits bacterial growth that follows a similar trajectory in the early phases, but at the transition to stationary phase there is a drop in culture density indicative of cell lysis and loss of viability (**Fig 1C,D**) [41]. Collectively, these results illustrate that ·NO alters PYO detection and that the co-occurrence of ·NO and PYO negatively impacts *P. aeruginosa* growth, suggesting that ·NO and PYO might be interacting.

### Nitric oxide reacts with phenazines, yielding chemically distinct products

Given the capacity of PYO in the presence of ·NO to alter *P. aeruginosa* growth, we proceeded to investigate the chemical interactions between PYO, ·NO, and phenazines in more detail. *P. aeruginosa* generally produces and secretes four structurally distinct phenazine metabolites that promote its anaerobic survival and are antagonistic against other organisms, including other microbes and humans, and thus can serve as natural antibiotics [42, 43]. We found that when PYO and the structurally similar 1-OHPhz are exposed to ·NO a color change occurs (**Fig 2A**) and a new product is formed (**Table 1**). These changes occur in both neutral and acidic conditions (**Fig 2A, Fig S2A,B**). Spectral analyses after ·NO exposure also suggest absorption properties consistent with phenazine metabolites, most notably an absorbance peak between 300 and 400 nm that is also present in the parent phenazine molecules (**Fig S2B,C**).

**Figure 2.**
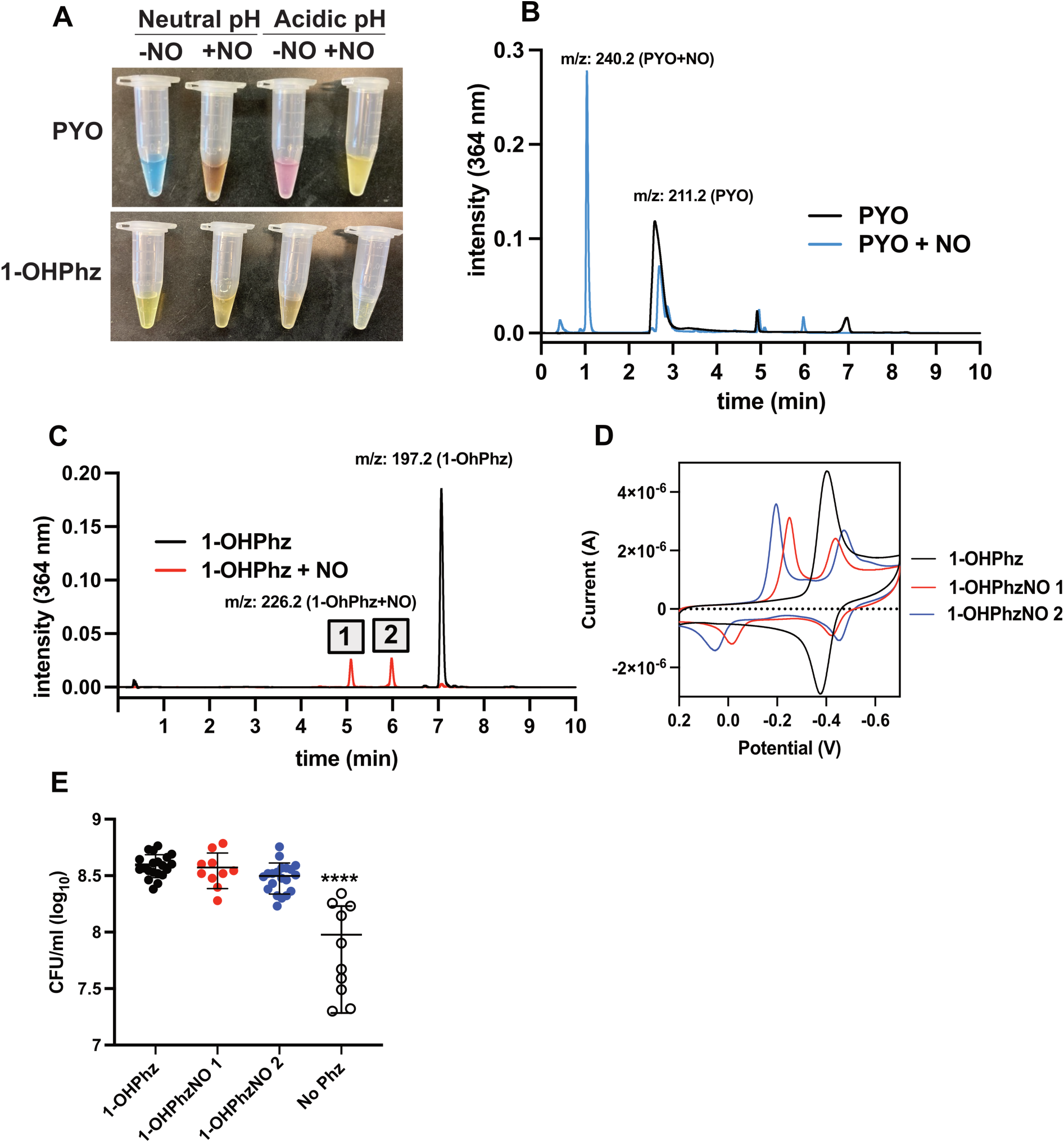
Nitric oxide reacts with phenazines to yield chemically distinct redox-active metabolites. **A)** Incubation of pyocyanin (PYO) or 1-hydroxyphenazine (1-OHPhz) at pH 7 or pH 4 in the presence of nitric oxide (NO). **B)** Chromatogram for absorbance at 364 nm and MS analyses for phenazines: m/z of PYO reacted with NO (PYO-NO) with reported. **C)** LC-MS analysis and m/z of 1-OHPhz reacted with ·NO (1-OHPhzNO) with absorbance at 364 nm reported and two products labeled #1 and #2. **D)** Cyclic voltammetry of 1-OHPhz and derivatives. **E)** Anaerobic survival after 24 h measured by colony forming units (CFU) per ml of *P. aeruginosa* provided with 1-OHPhz or its derivatives. ****p < 0.0001, one-way ANOVA with Tukey multiple comparisons vs ‘No Phz;’ n = 10-20 from 3 biological replicates, bars are mean +/− SD.

**Table 1.** Rf values for phenazine reactivity. Mobile Phase 1: 50:50 chloroform:methanol; Mobile Phase 2: 95:5 chloroform:methanol.

| Compound | Rf | Mobile Phase |
| --- | --- | --- |
| Pyocyanin | 0.79 | 1 |
| Pyocyanin-NO (acidic) | 0.44 | 1 |
| Pyocyanin-NO (neutral) | 0.45 | 1 |
| 1-OHPhz | 0.86 | 2 |
| 1-OHPhzNO 1,2 (acidic) | 0.55 | 2 |
| 1-OHPhzNO 1,4 (acidic) | 0.36 | 2 |
| 1-OHPhzNO 1,2 (neutral) | 0.55 | 2 |
| 1-OHPhzNO 1,4 (neutral) | 0.35 | 2 |

To determine the chemical nature of these ·NO-exposed phenazines, samples were analyzed using LC-MS, and mass to charge ratios (*m/z*) were compared to the parent molecule. PYO (*m/z* [M+H]^+^ = 211.2) possesses a retention time of ∼3 min, while a new product emerges following exposure to ·NO with a retention time of 1 min and mass addition of 29 (*m/z* 240.2) (**Fig 2B**). 1-OHPhz (*m/z* 197.2) exposed to ·NO displays two new masses (*m/z* 226.2) with distinct retention times and a mass addition of 29 Da relative to the parent molecule (**Fig 2C**). These products form over various pH ranges and in both the presence and absence of ambient oxygen (**Fig S2C,D; Table 1**). The chemical similarities between the PYO and 1-OHPhz ·NO-reacted products suggest a conserved reaction mechanism that might be revealed with additional experimentation with the two 1-OHPhz products (**Fig 2C, #1 and #2**). These products were subsequently purified and analyzed using cyclic voltammetry to assess their redox properties. Consistent with previous studies [23, 44], 1-OHPhz exhibits a reversible two-electron transfer redox reaction, characterized by a pair of symmetrical oxidation and reduction peaks with a midpoint potential of −390.5 mV vs Ag/AgCl and a peak potential separation of ∼28 mV (**Fig 2D**). By contrast, the ·NO-reacted 1-OHPhz products display more complex redox behavior: each product shows two pairs of oxidation and reduction peaks, with Product 1 showing midpoint potentials of −132 mV and −429 mV, and Product 2 showing midpoint potentials of −70 mV and −462.5 mV vs Ag/AgCl). These results suggest that they may undergo multistep redox processes or contain more than one redox-active center [45]. A qualitative assessment of biologically-mediated reduction of PYO and ·NO-reacted PYO via incubation with *P. aeruginosa* also suggests that these metabolites retain their redox activity (**Fig S2E**).

To assess the functional impact of this redox activity, we asked whether these modified phenazines could support anaerobic survival for *P. aeruginosa*. Cells were incubated anaerobically with an oxidizing potential +/− phenazine, and cellular survival was monitored over 24 h. In line with previous experiments [23], 1-OHPhz supports anaerobic survival (**Fig 2E**). Both ·NO-transformed phenazines also support anaerobic survival (**Fig 2E**), which is consistent with their maintenance of redox activity that is a required feature for this phenotype [24]. These results demonstrate that ·NO can transform phenazine metabolites and change their chemical properties, without disrupting their usage as electron acceptors for anaerobic survival.

The mass addition of 29 Da to both PYO and 1-OHPhz suggest a conserved reaction mechanism with NO (**Fig 3**). To determine the structure of ·NO-transformed phenazines, we analyzed the two purified 1-OHPhz products using ^1^H nuclear magnetic resonance (NMR) spectroscopy. The resulting spectra reveal the formation of an oxime group on the base pyrazine structure (**Fig 3**), which is consistent with FTIR results (**Fig S3**). To elucidate the mechanistic origins of the two possible oxime products, dispersion-corrected density functional theory calculations were employed using unrestricted dispersion-corrected DFT (μB97X-D/set cc-pvDZ-CPCM (H_2_O); see **Supporting Information** for additional **Computational Details** and **Figs S5-S11, Table S1**). ·NO is recognized as a stable free radical, reacting rapidly with redox active metals and other free radicals, yet relatively unreactive in processes involving two-electron oxidations or reductions [46]. However, the ability of ·NO to form N_2_O_2_, a weak electrophilic dimer, in the presence of aromatic hosts, lends itself to nucleophilic attack via the more negatively charged sites on 1-OHPhz [47].

**Figure 3.**
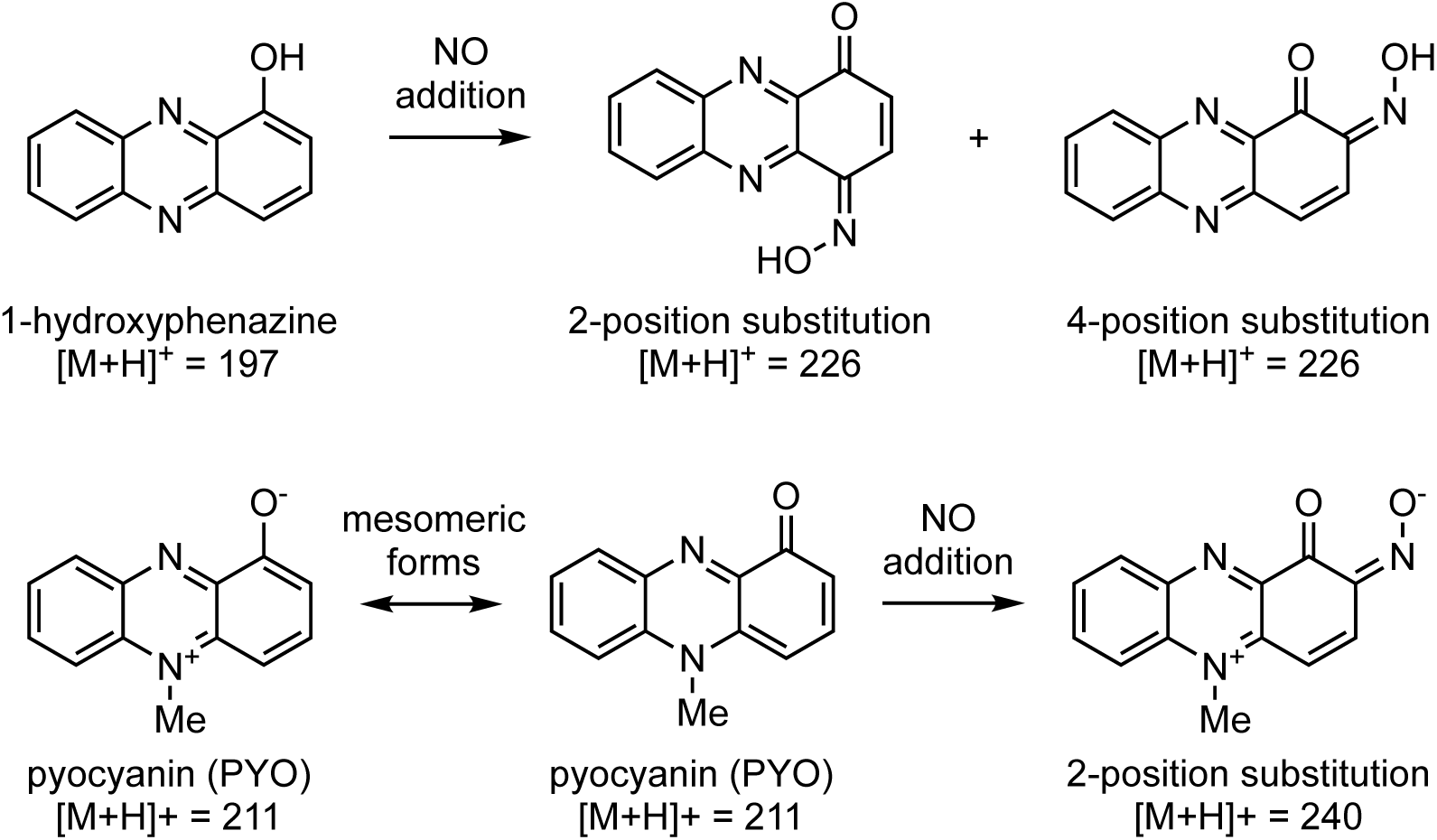
The chemical structures 1-hydroxyphenazine, pyocyanin, and their nitric oxide adducts, with their detected protonated masses shown.

In our case, N_2_O_2_ is introduced as the reactive partner by coordination with 1-OHPhz, approximately 3.10 Å above the aromatic rings (**Fig 4**). From this complex, we found that N_2_O_2_ then undergoes a nucleophilic attack (via a barrier of ∼21 kcal/mol) at either the two or four position due to their increased nucleophilicity (**Supporting Information**) leading to the corresponding NONO adducts. In turn, these adducts can undergo N-N homolytic cleavage to form ·NO and an open-shell hydroxyamino-like intermediate (^2^**3** and ^4^**3**) that can rapidly tautomerize (**Supporting Information**) to the lowest energy N-OH adducts [48]. Finally, both pathways undergo hydrogen atom transfer (HAT) via an energetically feasible barrier (∼26 kcal/mol) yielding the desired products.

**Figure 4.**
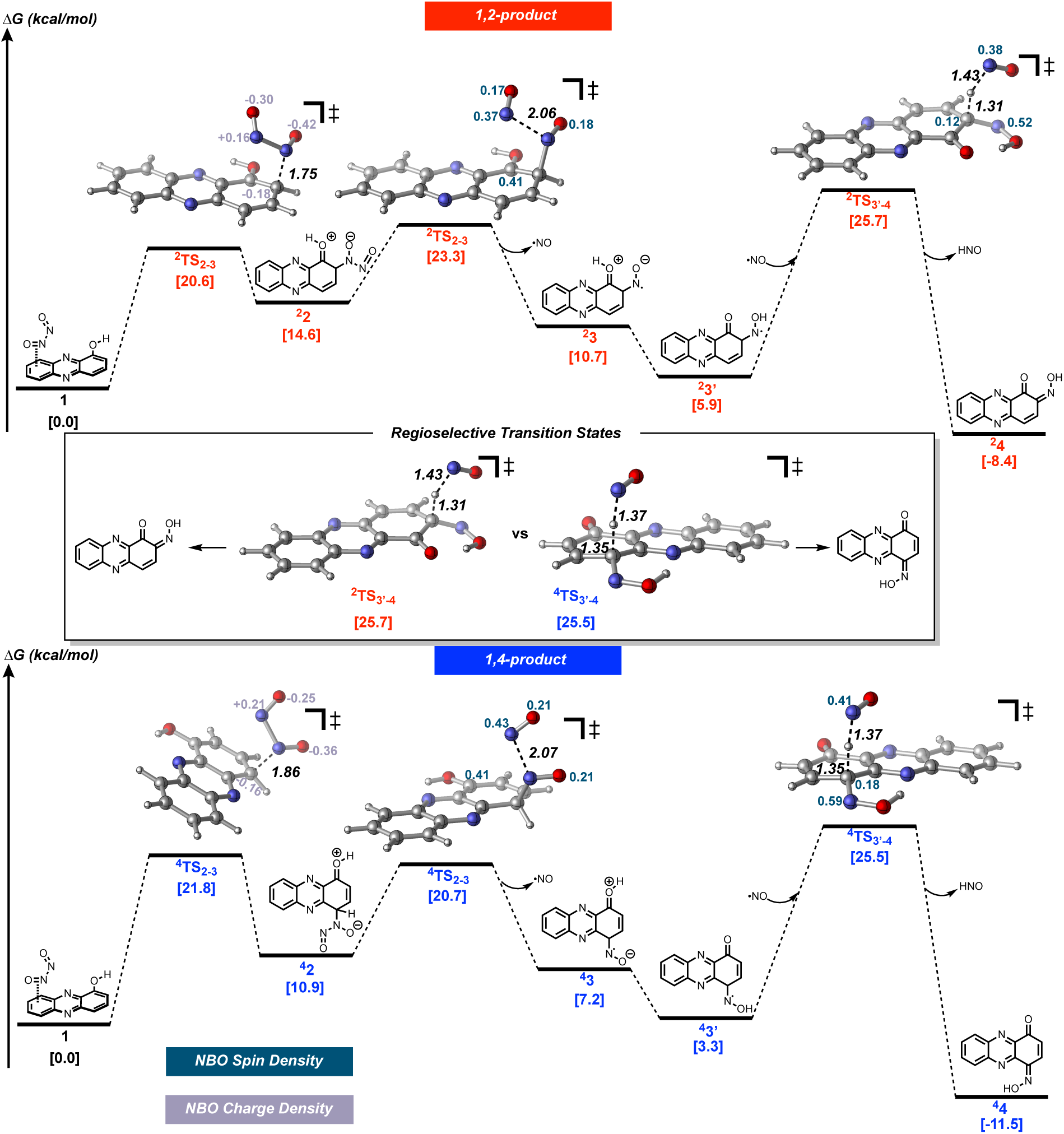
Proposed structures and reaction mechanisms of phenazine-NO interactions. Free energy pathway for the 1,2- and 1,4-addition of nitric oxide to 1-hydroxyphenazine. Free energies (in kcal/mol) were calculated at the uμb97xD/6-311+g(d,p)-CPCM(H_2_O)//uμb97xD/cc-pvDZ-cpcm(H_2_O) level of theory. Bond lengths are presented in angstroms, written in black.

Charge density analysis was key to determining the initial step of the reaction mechanism. Natural bond orbital (NBO) analysis was used to visualize the charge density at each atomic position. Initially an aza-Michael Addition type mechanism was postulated, but NBO calculations revealed that the 2- and 4-positions are electron rich. The charge densities for the N_2_O_2_ addition are shown in **Fig 4**, demonstrating the respective nucleophilicity and electrophilicity of each reactant. NBO spin density analysis was also used to confirm the open-shell nature of the homolytic cleavage and HAT transition states. The radical character is delocalized throughout the phenazine ring as seen by the selected spin densities (**Fig 4**). Due to the irreversibility of the HAT transition state, this was determined to be the regioselective step that establishes the structurally distinct products. All steps leading up to the HAT can be considered reversible and due to a difference of only 0.2 kcal/mol in the ΔG^‡^ of the regioselective transition states, a mixture of products is expected.

Taken together, these results show that ·NO chemically transforms some phenazine metabolites into two structurally distinct molecules via a mechanism that employs nucleophilic attack of the aromatic ring on the ·NO dimer, homolytic cleavage, and hydrogen atom transfer. While these nitrosylated phenazines maintain redox properties that can support *P. aeruginosa* anaerobic survival once they are formed, why the presence of PYO and ·NO negatively impacted *P. aeruginosa* during growth remained a puzzle that we next sought to address.

### PYO-NO reactivity is acutely toxic

To gain insight into why reaction between phenazines and ·NO are toxic to *P. aeruginosa*, we exposed *P. aeruginosa* to either ·NO, phenazines, or a combination of both for various periods to assess how their interactions affect cell viability. When *P. aeruginosa* is exposed acutely to high concentrations of ·NO for one hour, it loses viability (**Fig 5A**). When it is exposed to ·NO alongside 1-OHPhz or phenazine 1-carboxylic acid (PCA), there is a minor combined loss in viability (**Fig 5A**). Co-exposure to NO and phenazine 1-carboxamide (PCN) results in more pronounced death (**Fig 5A**). However, the combined toxicity for each of these phenazines is relatively minor compared to that effected by PYO and ·NO (**Fig 5A**), where ∼1-million-fold loss in viability is observed. To understand the kinetics of PYO-NO combined toxicity, cells were treated with propidium iodide (PI) during exposure to the molecules. PI is occluded from sufficiently polarized cellular envelopes but can diffuse across compromised envelopes, indicating these cells have a low metabolic activity and/or are dead [49, 50]. When cells are treated with either PYO or ·NO, there is no change in the PI relative fluorescence, which is similar to background (**Fig 5B**). However, when cells are incubated with PYO and ·NO simultaneously, there is a significant increase in PI relative fluorescence (**Fig 5B**). These results indicate that PYO reaction with ·NO causes rapid membrane depolarization, promoting PI cellular influx and suggesting a mechanism for cell death.

**Figure 5.**
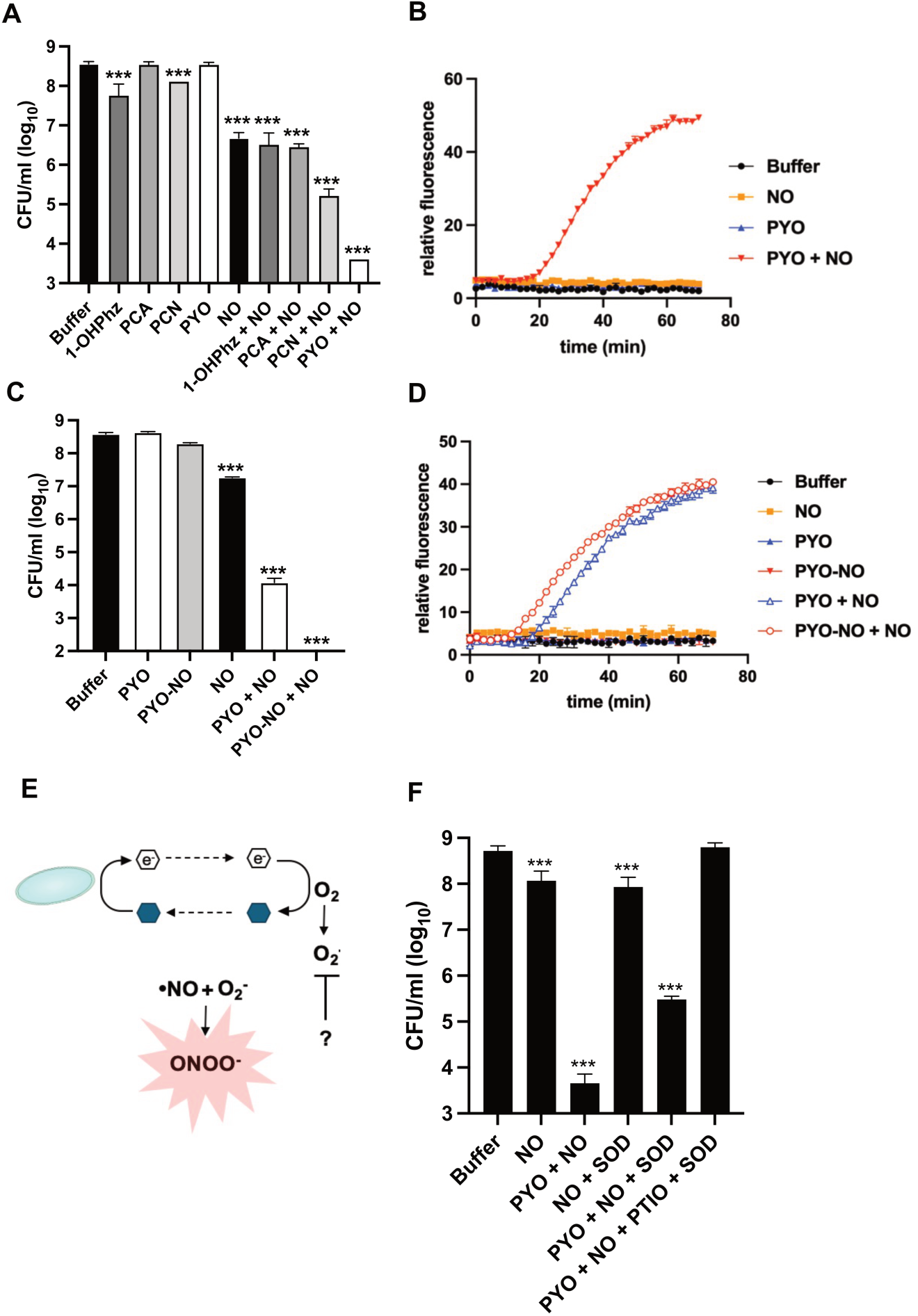
Pyocyanin-nitric oxide reactivity is acutely toxic to *P. aeruginosa*. **A)** Bacterial survival reported as CFU/mL of *P. aeruginosa* after incubation with 1-OHPhz, phenazine-1-carboxylic acid (PCA), phenazine-1-carboxamide (PCN), or PYO +/− ·NO; n = 3 biological replicates, mean +/− SD. **B)** Representative propidium iodide (PI) fluorescence of *P. aeruginosa* during incubation with PYO, NO, or PYO + ·NO. **C)** Bacterial survival reported as CFU/ml of *P. aeruginosa* after incubation with PYO, NO, PYO+NO or PYO-NO (product) + NO for one hour; n = 3 biological replicates, mean +/− SD. **D)** Representative PI fluorescence of *P. aeruginosa* during incubation with PYO, NO, or PYO + ·NO, or PYO-NO + ·NO. **E)** Proposed scheme for how PYO and ·NO may integrate with phenazine redox cycling and oxygen to produce the reactive nitrogen species ONOO^-^. **F)** Bacterial survival reported as CFU/mL of *P. aeruginosa* after incubation with PYO +/− ·NO +/− recombinant superoxide dismutase (SOD) +/− ·NO scavenger carboxy PTIO (PTIO) for one hour; n = 3 biological replicates, mean +/− SD. *** p < 0.001, one-way ANOVA with Tukey multiple comparisons vs buffer.

We wondered whether the product formed when PYO and ·NO react was causing the toxicity, or if the toxicity stemmed from how PYO reacts with NO. To distinguish between these possibilities, we exposed cells to ·NO, PYO, or the ·NO-derivative PYO-NO singly and in combination with ·NO and monitored viability. Incubation with each phenazine on its own did not significantly decrease cell viability (**Fig 5C**). Incubation with ·NO alone decreased viability ∼1log_10_ (**Fig 5C**). Incubation with PYO and ·NO simultaneously decreased viability by ∼4.5log_10_. Surprisingly, incubation of previously NO-reacted PYO (PYO-NO) with ·NO for a second time resulted in >8log_10_ loss in viability, with viability being below the limit of detection of 100 cells/mL (**Fig 5C**). These findings are consistent with the rapid increase in PI staining associated with PYO-NO reactivity with NO (**Fig 5D**). These results demonstrate that PYO-NO itself does not cause acute toxicity but that the toxicity derives from its formation and/or from subsequent reactions between PYO-NO and ·NO. The rapid depolarization of the bacterial cell envelope during these reactions rationalizes its efficient killing.

An important feature of phenazines produced by *P. aeruginosa* is their redox cycling capacity, which promotes *P. aeruginosa* anaerobic survival [23]. While such recycling can be achieved under strictly anoxic conditions using electrodes to regenerate oxidized phenazines, under (hyp)oxic conditions, reduced phenazines can be re-oxidized by molecular oxygen (O_2_) (**Fig 5E**). Yet phenazine oxidation by O_2_ produces superoxide radicals [51], which can then react with ·NO to produce peroxynitrite, one of the strongest nitrosative stress agents [52] (**Fig 5E**). Accordingly, we hypothesized that superoxide production may drive the rapid toxicity that occurs when PYO reacts with ·NO and reasoned that preventing superoxide formation might curtail this toxicity (**Fig 5E**). To test this hypothesis, we incubated cells with a combination of PYO and ·NO plus recombinant, exogenous superoxide dismutase (SOD), which catalyzes the dismutation of superoxide to hydrogen peroxide, thereby lowering superoxide concentration [53]. Cells were incubated for 1 hour with the various treatments and viability was determined. The addition of SOD had no effect on ·NO toxicity alone, but it did partially rescue the toxicity of PYO and ·NO provided simultaneously (**Fig 5F**). Addition of the ·NO scavenger carboxy-PTIO (PTIO) completely alleviates NO toxicity as well as PYO and ·NO combined toxicity (**Fig 5F**). However, ·NO scavenging must occur at the beginning of the incubation, since addition of PTIO after initial incubation with PYO and ·NO does not rescue *P. aeruginosa* viability (**Fig S4**). Together, these data demonstrate that ·NO can react with phenazines and impact cell viability depending on the ·NO-molecule pairing; further, the production of superoxide via redox cycling under oxic conditions contributes to this toxicity.

### Nitric oxide-enhanced PYO killing is species-specific and cell-intrinsic

Having established the likely mechanism whereby nitrosylation of PYO kills *P. aeruginosa*, we were curious whether other cell types were similarly sensitive. Strains of *Escherichia coli*, *Staphylococcus aureus*, and *Bacillus subtilis* were exposed to ·NO + PYO. When compared to *P. aeruginosa*, *E. coli* was more sensitive to PYO and NO cotreatment (**Fig 6A**). By contrast, both *S. aureus* and *B. subtilis* were more resistant to PYO and NO cotreatment compared to *P. aeruginosa*, with *S. aureus* being completely resistant (**Fig 6A**). The resistance of *S. aureus* to PYO and ·NO cotreatment was particularly interesting, since PYO has known antimicrobial effects against *S. aureus* [43]. To determine if the transformation of PYO by ·NO altered its antimicrobial activity, *S. aureus* was placed on a solid medium with a saturated disk of PYO or PYO-NO, and growth inhibition in the vicinity of the disk was measured. Consistent with previous results [43, 54], PYO inhibits *S. aureus* growth, but PYO-NO does not (**Fig 6C**). *B. subtilis*, *Acinetobacter baumannii*, and *E. coli* were also completely unphased by PYO-NO (**Fig 6C**). Because *S. aureus* frequently competes with *P. aeruginosa* for infectious niches, we wondered whether its resistance to PYO-NO might give it an advantage over *P. aeruginosa* in co-culture. To put this to the test, we compared the sensitivity of *S. aureus* and *P. aeruginosa* to PYO, ·NO, and the combination in mono or mixed cultures. While *P. aeruginosa* typically outcompetes *S. aureus* over time [55, 56], this one-hour incubation did not change the viability of *S. aureus* or *P. aeruginosa* (**Fig 6D**). Additionally, the exposure time to PYO was insufficient to decrease the viability of *S. aureus* (**Fig 6D**). By contrast, when *P. aeruginosa* was exposed to PYO and ·NO simultaneously, there is a 5log_10_ drop in viability, regardless of whether *S. aureus* is present (**Fig 6E**). Yet *S. aureus* remained completely resistant to PYO and ·NO simultaneous exposure (**Fig 6D**). These results illustrate that *S. aureus* possesses intrinsic resistance to PYO and ·NO reactivity. Collectively, these results demonstrate that PYO and ·NO toxicity is species specific.

**Figure 6.**
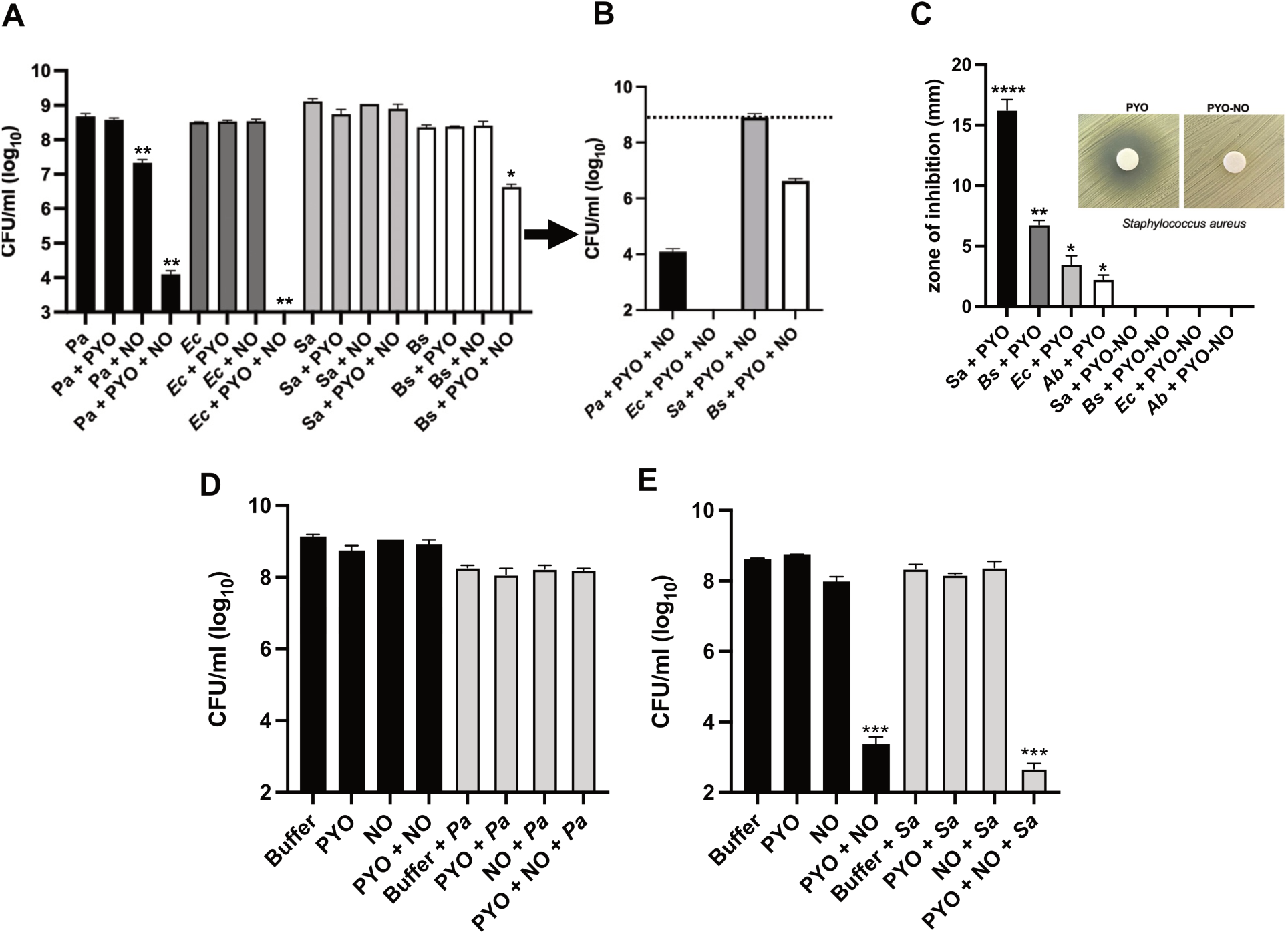
Pyocyanin-nitric oxide toxicity has species specificity with cell-intrinsic resistances. **A)** Bacterial survival reported as CFU/mL of *P. aeruginosa, Escherichia coli*, *Staphylococcus aureus*, and *Bacillus subtilis* after incubation with PYO, ·NO or PYO+NO for one hour, with **B)** inset to the right showing PYO+NO killing for each species; n=3 biological replicates, mean +/− SD. *p < 0.05, ***p < 0.001, one-way ANOVA with Tukey multiple comparison vs strain-only control. **C)** Radial zone of inhibition measurements for *B. subtilis*, *E. coli*, and *A. baumannii* after disk exposure for 24h to PYO or PYO-NO, with *S. aureus* insert; n = 3 biological replicates, mean +-SD. *p < 0.05, **p < 0.01, ****p < 0.0001, unpaired t-test PYO vs PYO-NO. **D)** Bacterial survival reported as CFU/ml of *S. aureus* +/− incubation with *P. aeruginosa* +/− PYO+NO; n = 3 biological replicates, mean +/− SD. **E)** Bacterial survival reported as CFU/ml of *P. aeruginosa* +/− incubation with *S. aureus* +/− PYO +/− NO; n = 3 biological replicates, mean +/− SD. ***p < 0.001, one-way ANOVA with Tukey multiple comparisons vs buffer.

## Discussion

·NO is commonly assumed to be broadly reactive, but specific molecular features react preferentially in ways that have important biological consequences. For example, the phenolic ring of tyrosine and the thiol side chain of cysteine are both reactive with ·NO and its derivatives, thereby altering protein function [57, 58]. The main contribution of this work is to expand this principle from protein biochemistry into the secreted small molecule landscape. By using nitrosylation of phenazine metabolites as a model system, we demonstrated that certain phenazine metabolites possessing 1-hydroxy functional groups are ·NO-reactive, and that this reactivity transforms the properties of the phenazines in a manner that alters their biological effects.

How does this change in impact manifest? Our results indicate that the change in reactivity of the nitrosylated phenazine and/or the formation of the adduct itself is responsible for its altered biological effects, but varies according to the specific phenazine and bacterial species in question. Specifically, we found that when PYO reacts with ·NO, it induces rapid loss of *P. aeruginosa* viability, but this loss does not occur when the structurally similar phenazine 1-OHPhz reacts with ·NO. Further, ·NO-mediated toxicity depends on the species that is in proximity to the metabolite-NO reaction. For instance, the Gram-positive pathogen *S. aureus* is intrinsically resistant to PYO reacting with ·NO and is also insensitive to the PYO-NO adduct. While the specific resistance mechanism awaits future determination, previous studies indicate *S. aureus* resists PYO and ·NO stress in part by using a lactic acid fermentative metabolism and inducing stress response and NO detoxification pathways [32, 54, 59, 60].

Both PYO and the structurally similar phenazine 1-OHPhz form new products when nitrosylated. Our structural analyses of 1-OHPhz reacted with ·NO identified both a 1,2-addition and 1,4-addition to the ring, while PYO reacted with ·NO results in only one product with the correct mass for ·NO addition. Considering the structural similarities between PYO and 1-OHPhz, the 1-hydroxy functional group is likely the conserved structural feature driving ·NO reactivity for both molecules, with the pyrazine methyl group likely preventing 1,4-addition to PYO. While we were able to detect PYO-NO reaction products in both oxic and anoxic environments, we found that the addition of PYO and ·NO simultaneously to growing cultures *P. aeruginosa* only results in cell death once the cultures reach stationary phase. An important change that occurs as *P. aeruginosa* transitions into this phase is increased hypoxia due to rapid oxygen consumption in a turbid culture, which can result in the chemical reduction of phenazines even when the air-liquid interface is oxic [61]. We thus infer that reduced PYO is preferentially reactive with ·NO and drives the rapid loss of viability over short time periods.

Phenazines are bacterial-derived secreted metabolites that are widespread in nature and disease, where they can support anaerobic metabolism and exert antimicrobial activity, among other functions [23, 27, 43, 62, 63]. The environments in which they are made are also replete with ·NO, which can be biotically or abiotically-derived [64–67]. The widespread occurrence of ·NO in small molecule-dominated ecosystems suggests that ·NO-mediated small molecule transformation may be an important and overlooked chemical process. Consistent with this prediction, a recent analysis of wastewater identified an ·NO-modified version of ciprofloxacin, which is a synthetic antibiotic with important clinical relevance [68]. Given the importance of both secreted metabolites and ·NO for both human and microbial physiology, our findings suggest that these ·NO-mediated transformations may have important implications for human health and the environment.

A major challenge in understanding small molecule function is dissecting their roles in complex ecosystems. This problem is compounded by our limited capacity to identify unknown metabolites from chemically complex environments without *a priori* knowledge [69]. Our analysis of ·NO-transformed phenazines illustrates that specific mass and polarity shifts occur following ·NO-phenazine reactivity, which suggests these chemical changes could be used to identify other molecules undergoing ·NO-mediated transformations. Here we have taken a reductionist approach to identify chemical parameters that facilitate ·NO reactivity with phenazines and optimized analytic methods to detect them while defining their biological importance; we hope to apply this logic to diverse and dynamic ecosystems with the goal of uncovering the breadth of metabolite chemical transformations facilitated by ·NO and their consequences.

## Materials and Methods

### Bacterial Strains and Reagents

Experiments were performed using *Pseudomonas aeruginosa* strain UCBPP-PA14 (*Pa*) unless otherwise noted. Experiments involving liquid culture were performed in lysogeny broth (LB) at 37 °C with aeration at 250 rpm (Innova) unless otherwise stated. Nitric oxide donors DETA-NONOate (Cayman #82120) and DEA-NONOate (#82100) were acquired from Cayman Chemicals. Other strains utilized were *Staphylococcus aureus* USA300 LAC, *Acinetobacter baumannii* ATCC 17978, *Escherichia coli* strain MG1655, *Bacillus subtilis* strain 168, all from the Newman lab strain collection.

### Growth in Nitric Oxide

Growth in the presence of nitric oxide was performed in either 5 mL volumes in 15 mL culture tubes or in 96-well plates, and in both cases NO was provided via DETA-NONOate. Bacterial strains were grown overnight in LB, and the following day cultures were diluted to OD_500_ of 0.05 in fresh media. DETA-NONOate stocks (500 mM in 10 mM NaOH) were thawed from −80°C and diluted to indicated concentrations in LB before pipetting 150 μl into a 96-well flat bottom plate or mixing with 5 mL LB in a 15 mL culture tube. For PYO supplementation experiments, PYO was diluted into LB at 100 μM. Diluted bacteria were inoculated 1:30 into the 96-well plate or into 15 mL culture tube. 96-well plates were placed in a Biotek Epoch2 plate reader at 37°C with continuous orbital shaking and OD_500_ absorbance readings captured hourly, and 15 mL culture tubes were placed into a 37°C incubator with 250 rpm shaking, and samples were taken over time for absorbance readings.

### Thin Layer Chromatography

Phenazines were spotted on analytical silica TLC plates (vendor) using 5 mL capillaries and allowed to dry. Dried plates were placed in small vial with the following mobile phases: PYO and derivatives utilized a mobile phase of 50:50 methanol:chloroform, while 1-OHPhz and derivatives utilized a mobile phase of 5:95 methanol:chloroform. Samples were ran until solvent front reached near end of plate and marked, with Rf values calculated as a fraction a product migrated as a function of the solvent front.

### Disk Diffusion Assay

Overnight bacterial cultures were streaked onto LB agar using a sterile cotton-tipped applicator. Sterile paper disks were placed onto the plate and spotted with 10 ul of 2 mM compound or solvent control. Plates were placed at 37°C incubator overnight, and radial zone of inhibitions were measured.

### Pyocyanin Purification

Pyocyanin (PYO) was synthesized and purified as previously described [70]. Briefly, ∼500 mg of phenazine methosulfate (Sigma, CAS #299-11-6) was dissolved in 500 mL 20 mM ammonium bicarbonate and placed in a fluorescent light box overnight at room temperature and constant stirring. The following day, an aqueous extraction was performed 6x with 100 mL washes of dichloromethane (DCM; Baker) in a separatory funnel and fractions collected and pooled. DCM extract was then acidified with 125 mL 0.1 M HCl, and the aqueous phase was extracted twice. Extracted phases were mixed with 2.5 mL 10 M NaOH and extracted 6x with 100 mL DCM and fractions collected and pooled. DCM was removed via Rotovap, and resulting solid was resolubilized in ∼5 mL methanol:DCM (20:80). Samples was dried overnight under N_2_ gas stream. The following day, sample was dissolved in 2 mL DCM and washed with 30 mL *n*-hexanes. Material was collected on vacuum filter and transferred to permanent storage vial. Purity was assessed using LC-MS.

### Phenazine Reactivity and Extraction

For nitric oxide reactivity with pyocyanin (PYO) and 1-hydroxyphenazine (1-OHPhz), purified PYO or commercially-available 1-OhPhz (Fisher; CAS #528-71-2) was resuspended in 20 mM HCl. Phenazines were then either diluted to 500 μM in HCl and sparged with N_2_ gas for 10 min, or diluted to 2 mM with deoxygenated 20 mM HCl or 1X PBS within an anaerobic chamber (Coy Laboratory Products). For oxic conditions, phenazines were diluted in ambient air and ambient air-exposed buffers. For sparged samples, phenazines were exposed for NO gas (Mesagas; ID 1660) with a relative flow rate of 10 mL/min for up to 2 h. For samples in anaerobic chamber, each reaction volume, 6 μl of 500 mM DEA-NONOate in 10 mM NaOH, and reaction proceeded overnight. PYO nitroso products (PYO-NO) were extracted via salting out with MgSO_4_ (1 g) and 1 mL acetonitrile (ACN; Fisher). ACN fractions were separated via preparative thin layer chromatography (TLC) (Sigma) and separated with a mobile phase of 1:1 methanol:chloroform (Sigma). For 1-OHPhz, nitroso products (1-OHPhzNO) precipitated following reactivity and were resolubilized in methanol prior to separation via preparative TLC and a mobile phase of 5:95 methanol:chloroform. In both cases, desired products were eluted with methanol from silica via flash chromatography. Fractions were collected and dried under N_2_ stream, and purity was assessed using LC-MS.

### Cyclic Voltammetry

Cyclic voltammetry (CV) experiments were performed using a CH Instruments potentiostat (CHI760E) at a scan rate of 20 mV/s. The electrochemical cell consisted of a 3-mm glassy carbon working electrode (BASi), an Ag/AgCl (3M KCl) reference electrode (BASi), and a platinum wire counter electrode. Phenazines and phenazine-NO derivatives were prepared as 10 mM stock solutions in methanol and subsequently diluted 100-fold to a final concentration of 100 µM in 1x phosphate-buffered saline (PBS, pH 7.3). All solutions were purged with N_2_ for at least 10 minutes prior to measurement, and experiments were conducted under a continuous N_2_ flow. Before each run, the glassy carbon electrode was polished using 0.05 μm gamma alumina powder.

### Anaerobic Survival Assay

Anaerobic survival experiments were performed using a 96-potentiostat system as previously described [71]. Cells were cultured for 20 hours in a *Pseudomonas* MOPS minimal medium (100 mM MOPS pH 7.2, 43 mM NaCl, 93 mM NH_4_Cl, 3.7 mM KH_2_PO_4_, 1 mM MgSO_4_), pelleted for 5 min at 8,000 *x g*, washed twice in minimal medium, resuspended at OD_500_ of 75, and transferred to anaerobic chamber with an ambient temperature of 33°C and ∼2% H_2_. Bacteria were diluted to OD of 15 with N_2_-sparged media, and 10 ul of bacteria were placed into a 96-well electrochemical plate containing 190 ul of media and 1-OHPhz or its derivative phenazines. The plate contained a carbon working electrode and counter electrode, plus a Ag/AgCl reference electrode. Plates were covered with a slit-seal cover (BioChromato #R80.120.00) and aluminum cover (Diversified Biochem #ALUM-1000). Plates were incubated statically at 33°C in a 96-potentiostat system, with each well held at 0 mV vs. Ag/AgCl reference electrode. Cyclic voltammograms and current readouts were collected over 24 hours, at which point 10 ul of cells was removed for CFU plating. All plastics for use within the anaerobic chamber were present for at least three days before use to allow sufficient degassing.

### Structural Analyses

#### Infrared Spectroscopy

Purified phenazines were analyzed via Fourier Transform Infrared (FTIR) spectroscopy on a Nicolet 6700 spectrometer. Solid samples were loaded onto the spectrometer fitted with a diamond-tipped ATR adaptor under a continuous N_2_ stream at a flow rate of 40ft^3^/hr. Spectra were captured for 64 scans with Resolution 4. Corrected interferogram was obtained by subtracting a scan background blank, and spectral values were exported from the OMNIC software subsequent plotting.

#### Nuclear Magnetic Resonance

##### Compound Preparation

Phenazines were reacted and extracted in bulk as described in the “Phenazine Reactivity and Extraction” section. Approximately 5 mg of product was solubilized with deuterated methanol and loaded into 5 mm, 7 inch NMR tubes (Norell).

##### Chemical Methods

^1^H NMR spectra were recorded on Varian Inova spectrometer (500 MHz), in the stated solvents as a reference for the internal deuterium lock. The chemical shift data for each signal are given as δ_H_ in units of parts per million (ppm) relative to tetramethylsilane (TMS) where δ_H_ (TMS) = 0.00 ppm. The spectra are calibrated using the solvent peak with the data provided by Fulmer *et al* [72]. The multiplicity of each signal is indicated by s (singlet); br s (broad singlet); d (doublet); dd (doublet of doublets), ddd (doublet of doublet of doublets), t (triplet), q (quartet), dq (double of quartet) or m (multiplet). The number of protons (n) for a given resonance signal is indicated by nH. Where appropriate, coupling constants (*J*) are quoted in Hz and are recorded to the nearest 0.1 Hz. Identical proton coupling constants (*J*) are averaged in each spectrum and reported to the nearest 0.1 Hz. The coupling constants were determined by analysis using MestReNova version 14.3.0 software. Infrared (IR) spectra were obtained from neat samples, either as liquids or solids using a diamond ATR module. The spectra were recorded on a Nicolet 6700 spectrometer spectrometer. Absorption maxima are reported in wavenumbers (cm^−1^).

###### 2-(Hydroxyimino)phenazin-1(2H)-one

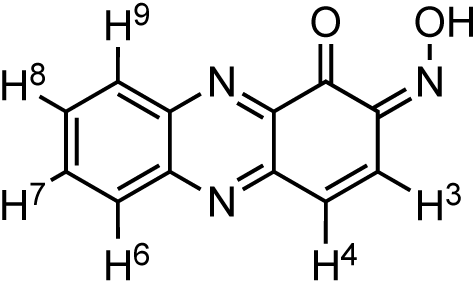

***v̅***_max_ (solid)/cm^−1^ 3309, 2925, 2854, 1742, 1608, 1509, 1251, 1177, 1037, 829; ^1^H NMR (500 MHz; MeOD) δ_H_ 8.33 (2H, d, *J* 8.2, H^7^ & H^8^), 8.28 (1H, d, *J* 10.4, H^4^), 8.02 (1H, dd, *J* 8.2, 7.7, H^9^), 7.95 (1H, dd, *J* 8.2, 7.7, H^6^), 6.78 (1H, d, *J* 10.4, H^3^); LRMS *m/z* (ESI^+^) 226.2 ([M+H]^+^ 100%)

###### 4-(Hydroxyimino)phenazin-1(4H)-one

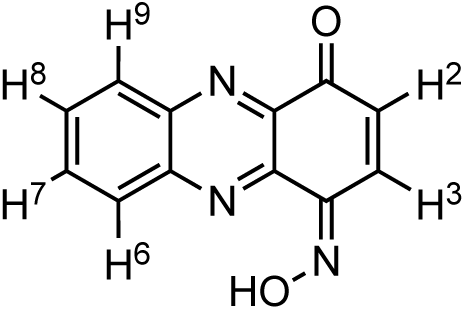

***v̅***_max_ (solid)/cm^−1^ 3309, 2925, 2854, 1742, 1608, 1509, 1251, 1177, 1037, 829; ^1^H NMR (500 MHz; MeOD) δ_H_ 8.38 (2H, d, *J* 8.6, H^6^), 8.14 (1H, d, *J* 8.2, H^9^), 7.98 (1H, ddd, *J* 8.2, 7.7, H^7^), 7.91 (1H, dd, *J* 8.0, 7.7, H^8^), 7.65 (1H, d, *J* 10.2, H^2^), 6.85 (1H, d, *J* 10.2, H^3^); LRMS *m/z* (ESI^+^) 226.2 ([M+H]^+^ 100%).

The above represents a doublet appearing at 4.56ppm on the corrected (to the residual non-deuterated solvent) x axis. This peak integrates to two protons and has a coupling constant (also referred to as J value) of 7.23Hz.

### Phenazine-Nitric Oxide Toxicity

Overnight bacterial cultures were pelleted for 3 min at 8000 *x* g at room temperature and resuspended at an OD_500_ of 1 for *Pa* in PBS. For anaerobic experiments, pellets were taken to anaerobic chamber and resuspended in de-gassed PBS. Cells were diluted to OD_500_ of 1 for *Pa*, or ∼10^8^ colony-forming units (CFU) per mL for other bacterial species. Cells were incubated for 1 h with 1.5 mM DEA NONOate (or indicated concentration) +/− 100 μM phenazine at room temperature and serial-diluted on LB agar for CFU enumeration. For bacterial interaction experiments, equal ratios of *P. aeruginosa* and *S. aureus* were mixed with a final OD_500_ of 1 and incubated with DEA NONOate +/− phenazine, and surviving cells were recovered by selective plating on mannitol salt agar to select for *S. aureus* or *Pseudomonas* isolation agar to select for *P. aeruginosa*.

### Propidium Iodide Spectroscopy

Overnight bacterial cultures were pelleted for 3 min at 8,000 x g and resuspended in an equal volume of 1X PBS. OD_500_ values were obtained and were normalized to OD_500_ = 1 in PBS. DEA-NONOate was provided as the NO donor at a final concentration of 1.5 mM, and PYO or PYO-NO was supplemented at a final concentration of 100 μM. Propidium iodide (PI) was added at a final concentration of 66.8 μM. After combining all reagents with cells, 100 μl of cell suspensions were placed in black-sided 96-well plates and loaded onto a SpectraMax M5 multi-mode microplate reader. PI fluorescence was monitored every minute (ex: 535, em: 620).

### LC-MS

Phenazine samples were analyzed as previously described [73, 74]. Briefly, samples were autosampled from 10°C into a Waters LC-MS system (Waters e2695 Separations Module, 2998 PDA Detector, QDA Detector) with 10 ul injections onto a reverse phase C-18 column (XBridge #186006035) with a running gradient of 98% H2O to 88% acetonitrile over 11 min (run times were 20 min total) with 2% methanol throughout. UV-Vis and positive MS scans were acquired for each run. PYO and 1-OHPhz were distinguished by retention time (∼3 min and ∼7 min, respectively), detected at 364 nm, and manually verified by examining masses 211.2 (PYO) and 197.2 (1-OhPhz). NO-reacted phenazines were detected at 364 nm and had predicted masses of 240.2 (PYO-NO) and 226.2 (1-OhPhz 1 & 2). Peaks were automatically mapped in the UV-Vis channels by retention time and the LC trace at 364 nm was exported as a text file from the Empower software.

### UV-VIS Spectroscopy

Phenazine stocks were diluted to 100 μM into either 1X PBS for neutral buffering or 20 mM HCl for acidic conditions. Samples were placed into a quartz 96 well plate and absorbance was measured between 200-800 nm with 5 nm steps and no shaking on a SpectraMax Multi-Mode microplate reader.

### Reaction Mechanism Calculations

All geometry optimizations of intermediates and transition states were achieved using the spin unrestricted uwb97xD [75] /cc-pvDZ [76] method, in water using the CPCM solvent model [77–81] as implemented in Gaussian16 [82]. All calculations used the “guess=mix,always” keywords and “opt=noeigen” was implemented for transition states. Frequency calculations were also conducted at the same level of theory to obtain vibrational frequencies to determine the identity of the stationary points as intermediates (no imaginary frequencies) or as transition states (only one imaginary frequency), as well as obtaining the thermochemistry: enthalpy (DH) and free energy (DG) at the temperature of 298 K. All spin and charge densities were done using the “pop=nbo” [83] keyword at the uwb97xD/6-311+g(d,p)-cpcm(H_2_O)//uwb97xD//cc-pvDZ-cpcm(H_2_O) level of theory. Extensive conformational analysis was performed using CREST [83–85] version 3.0.2 with XTB [86, 87] version 6.7.1 and only the lowest-energy species are shown and discussed. All structural figures were generated with CYLview [88]. Distances in structural figures are shown in Å and energies are in kcal/mol. Single Point energy corrections were carried out further with the following methods.

i. uwb97xD/6-311+g(d,p)-CPCM(H_2_O)//uwb97xD/cc-pvDZ-cpcm(H_2_O)
ii. uwb97xD/aug-cc-pvTZ-CPCM(H_2_O)// uwb97xD/cc-pvDZ-cpcm(H_2_O)
iii. uwb97xD/aug-cc-pvQZ-CPCM(H_2_O)// uwb97xD/cc-pvDZ-cpcm(H_2_O)

## Acknowledgements

We thank members of the Newman Laboratory for critical evaluation of this manuscript. This work was supported by Jane Coffin Childs Memorial Fund for Medical Research (ZRL), National Institutes of Health (NIH) K22AI182146 (ZRL), the Doren Family Foundation (DKN), NIH R01HL152190 (DKN), and NIH 2R01AI27850-06A1 (DKN). SJC is grateful to Michael and Alice Jung for endowing the Jung Chair in Medicinal Chemistry and Drug Discovery at UCLA, which partially supported this work. MS and OG were supported by NIH R35GM137797 MIRA grant.

## Notes

### Competing Interest Statement

The authors have declared no competing interest.

