## Supplemental Information for "Nitric oxide tunes secreted metabolite bioactivity"

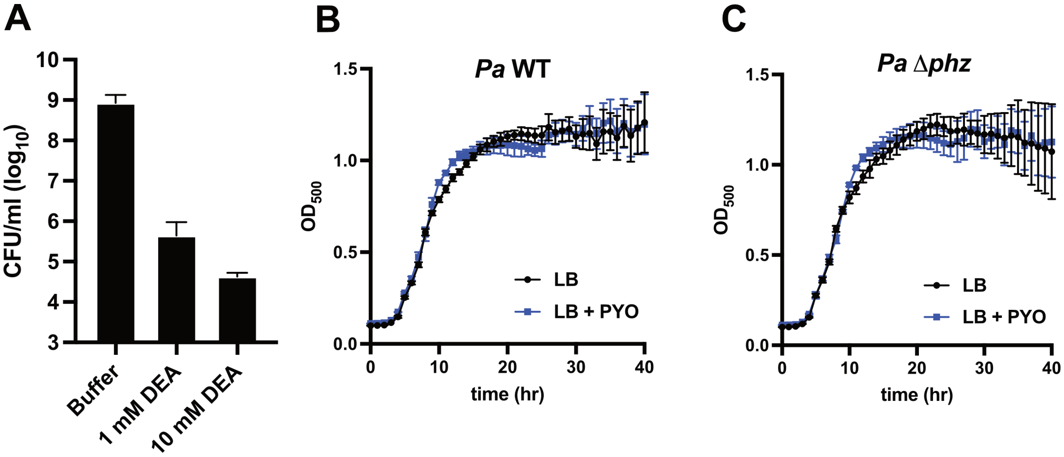


**Figure S1. Nitric oxide is acutely toxic to *P. aeruginosa*, and PYO does not negatively impact *Pa* growth. A)** CFU/mL after 1 h exposure to NO delivered via the small molecule donor DEA-NONOate; n = 2 biological replicates, mean +/- SD. **B)** Representative growth monitored by absorbance at 500 nm of wildtype (WT) *Pa* with 100 μM PYO supplemented at the start of growth; n = 3 technical replicates from 1 of 3 biological replicates, mean +/- SD. **C)** Growth monitored by absorbance at 500 nm of Δ*phz Pa* with 100 μM PYO supplemented at the start of growth; n = 3 technical replicates from 1 of 3 biological replicates, mean +/- SD.

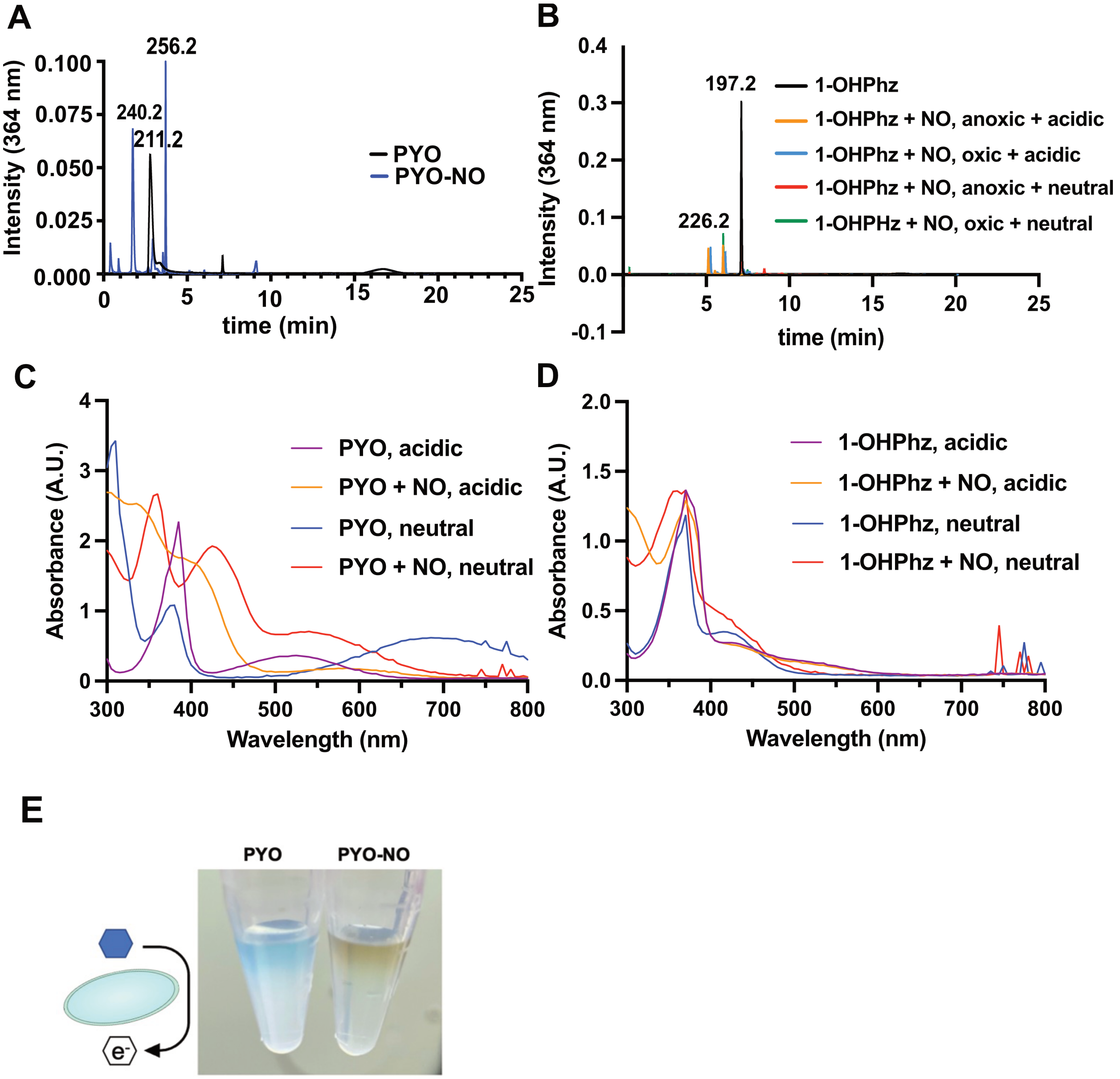


**Figure S2. Nitric oxide reacts with phenazines to yield chemically distinct redox-active metabolites. A)** Chromatogram at 364 nm with masses of PYO reacted with NO in oxic, acidic conditions indicated. **B)** Chromatogram at 364 nm of LC/MS analysis of 1-OHPhz reacted with NO in oxic and anoxic conditions, and at acidic or neutral pH. **C)** UV-Vis analysis of 100 µM PYO after NO reactions performed in acidic or neutral conditions. **D)** UV-Vis analysis of 100 µM 1-OHPhz after NO reactions performed in acidic or neutral conditions. **E)** Incubation of *Pa* with 100 µM PYO or PYO reacted with NO (PYO-NO).

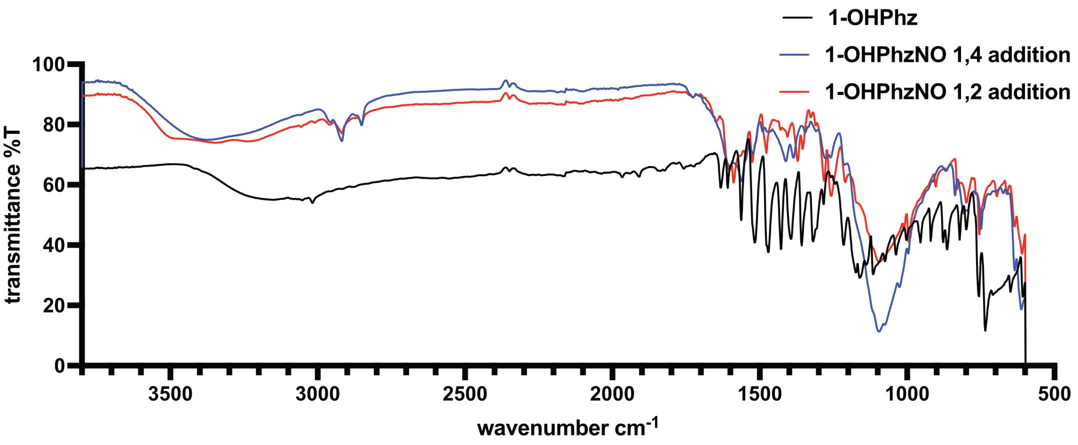


**Figure S3. Spectral analysis of 1-OHPhz following reactivity with NO.** FTIR analysis of 1-OHPhz and derivatives

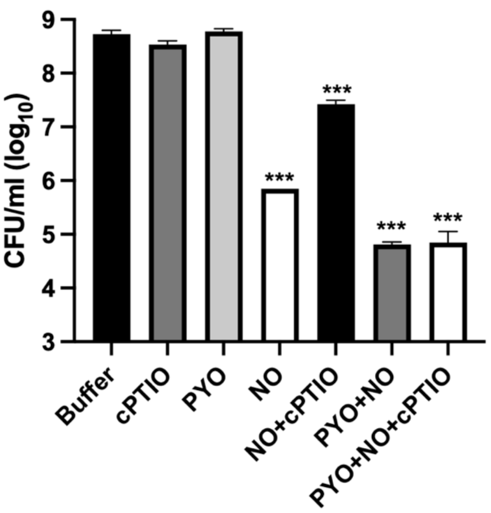


**Figure S4. PYO-NO reactivity is acutely toxic to *P. aeruginosa*. A)** Bacterial survival reported as CFU/mL of *Pa* after incubation with PYO +/- NO, with the addition of NO scavenger carboxy-PTIO (cPTIO) after 1 h of exposure; n = 2 biological replicates, mean +/- SD. ***$p < 0.001$, one-way ANOVA with Tukey multiple comparisons vs buffer.

### Supplementary Computational Details

All geometry optimizations of intermediates and transition states were achieved using the spin unrestricted uwb97xD [Ref1]/cc-pvDZ [Ref2] method, in water using the CPCM solvent model [Ref3] as implemented in Gaussian16 [Ref4]. All calculations used the "guess=mix,always" keywords and "opt=noeigen" was implemented for transition states. Frequency calculations were also conducted at the same level of theory to obtain vibrational frequencies to determine the identity of the stationary points as intermediates (no imaginary frequencies) or as transition states (only one imaginary frequency), as well as obtaining the thermochemistry: enthalpy (DH) and free energy (DG) at the temperature of 298 K. All spin and charge densities were done using the "pop=nbo" [Ref5] keyword at the uwb97xD/6-311+g(d,p)-cpcm($H_2O$)//uwb97xD//cc-pvDZ-cpcm($H_2O$) level of theory. Extensive conformational analysis was performed using CREST [Ref6] version 3.0.2 with XTB [Ref7] version 6.7.1 and only the lowest-energy species are shown and discussed. All structural figures were generated with CYLview [Ref8]. Distances in structural figures are shown in Å and energies are in kcal/mol. Single Point energy corrections were carried out further with the following methods.

i) uwb97xD/6-311+g(d,p)-CPCM($H_2O$)//uwb97xD/cc-pvDZ-cpcm($H_2O$)
ii) uwb97xD/aug-cc-pvTZ-CPCM($H_2O$)// uwb97xD/cc-pvDZ-cpcm($H_2O$)
iii) uwb97xD/aug-cc-pvQZ-CPCM($H_2O$)// uwb97xD/cc-pvDZ-cpcm($H_2O$)

1. (a) J.-D. Chai and M. Head-Gordon, "Systematic optimization of long-range corrected hybrid density functionals," *J. Chem. Phys.*, **128** (2008) 084106. (b) J.-D. Chai and M. Head-Gordon, "Long-range corrected hybrid density functionals with damped atom-atom dispersion corrections," *Phys. Chem. Chem. Phys.*, **10** (2008) 6615-20
2. T. H. Dunning Jr., "Gaussian basis sets for use in correlated molecular calculations. I. The atoms boron through neon and hydrogen," *J. Chem. Phys.*, **90** (1989) 1007-23
3. (a) Klamt, A.; Schüürmann, G. COSMO: a new approach to dielectric screening in solvents with explicit expressions for the screening energy and its gradient. *J. Chem. Soc. Perkin Trans. 2* **1993**, *0*, 799-805. (b) Tomasi, J.; Persico, M. Molecular Interactions in Solution: An Overview of Methods Based on Continuous Distributions of the Solvent. *Chem. Rev.* **1994**, *94*, 2027-2094. (c) Andzelm, J.; Kölmel, C.; Klamt, A. Incorporation of solvent effects into density functional calculations of molecular energies and geometries. *J. Chem. Phys.* **1995**, *103*, 9312-9320. (d) Barone, V.; Cossi, M. Quantum Calculation of Molecular Energies and Energy Gradients in Solution by a Conductor Solvent Model. *J. Phys. Chem. A* **1998**, *102*, 1995-2001. (e) Cossi, M.; Rega, N.; Scalmani, G.; Barone, V. Energies, structures, and electronic properties of molecules in solution with the C-PCM solvation model. *J. Comput. Chem.* **2003**, *24*, 669-681.
4. Gaussian 16, Revision C.01, Frisch, M. J.; Trucks, G. W.; Schlegel, H. B.; Scuseria, G. E.; Robb, M. A.; Cheeseman, J. R.; Scalmani, G.; Barone, V.; Petersson, G. A.; Nakatsuji, H.; Li, X.; Caricato, M.; Marenich, A. V.; Bloino, J.; Janesko, B. G.; Gomperts, R.; Mennucci, B.; Hratchian, H. P.; Ortiz, J. V.; Izmaylov, A. F.; Sonnenberg, J. L.; Williams-Young, D.; Ding, F.; Lipparini, F.; Egidi, F.; Goings, J.; Peng, B.; Petrone, A.; Henderson, T.; Ranasinghe, D.; Zakrzewski, V. G.; Gao, J.; Rega, N.; Zheng, G.; Liang, W.; Hada, M.; Ehara, M.; Toyota, K.; Fukuda, R.; Hasegawa, J.; Ishida, M.; Nakajima, T.; Honda, Y.; Kitao, O.; Nakai, H.; Vreven, T.; Throssell, K.; Montgomery, J. A., Jr.; Peralta, J. E.; Ogliaro, F.; Bearpark, M. J.; Heyd, J. J.; Brothers, E. N.; Kudin, K. N.; Staroverov, V. N.; Keith, T. A.; Kobayashi, R.; Normand, J.; Raghavachari, K.; Rendell, A. P.; Burant, J. C.; Iyengar, S. S.; Tomasi, J.; Cossi, M.; Millam, J. M.; Klene, M.;

Adamo, C.; Cammi, R.; Ochterski, J. W.; Martin, R. L.; Morokuma, K.; Farkas, O.; Foresman, J. B.; Fox, D. J. Gaussian, Inc., Wallingford CT, 2016.
5. J. P. Foster and F. Weinhold, "Natural hybrid orbitals," *J. Am. Chem. Soc.*, 102 (1980) 7211-18
6. (a) Pracht, P.; Bohle, F.; Grimme, S.; *Automated exploration of the low-energy chemical space with fast quantum chemical methods*, *Phys. Chem. Chem. Phys.*, **2020**, *22*, 7169-7192. (b) Pracht, P.; Grimme, S.; Bannwarth, C.; Bohle, F.; Ehlert, S.; Feldmann, G.; Gorges, J.; Müller, M.; Neudecker, T.; Plett, C.; Spicher, S.; Steinbach, P.; Wesołowski, P.A.; Zeller, F.; *CREST — A program for the exploration of low-energy molecular chemical space*, *J. Chem. Phys.*, **2024**, *160*, 114110. (c) Grimme, S.; *Exploration of Chemical Compound, Conformer, and Reaction Space with Meta-Dynamics Simulations Based on Tight-Binding Quantum Chemical Calculations*, *J. Chem. Theory Comput.*, **2019**, *15 (5)*, 2847-2862.
7. (a) Grimme, S.; Bannwarth, C.; Shushkov, P.; *A Robust and Accurate Tight-Binding Quantum Chemical Method for Structures, Vibrational Frequencies, and Noncovalent Interactions of Large Molecular Systems Parameterized for All spd-Block Elements (Z = 1-86)*. *J. Chem. Theory Comput.*, **2017**, *13 (5)*, 1989-2009. (b) Bannwarth, C.; Ehlert, S.; Grimme, S.; *GFN2-xTB — An Accurate and Broadly Parametrized Self-Consistent Tight-Binding Quantum Chemical Method with Multipole Electrostatics and Density-Dependent Dispersion Contributions J. Chem. Theory Comput.* **2019**, *15 (3)*, 1652–1671.
8. Legault, C. Y. (2009) CYLview, 1.0b, Université de Sherbrooke: Sherbrooke, Canada, http://www.cylview.org.

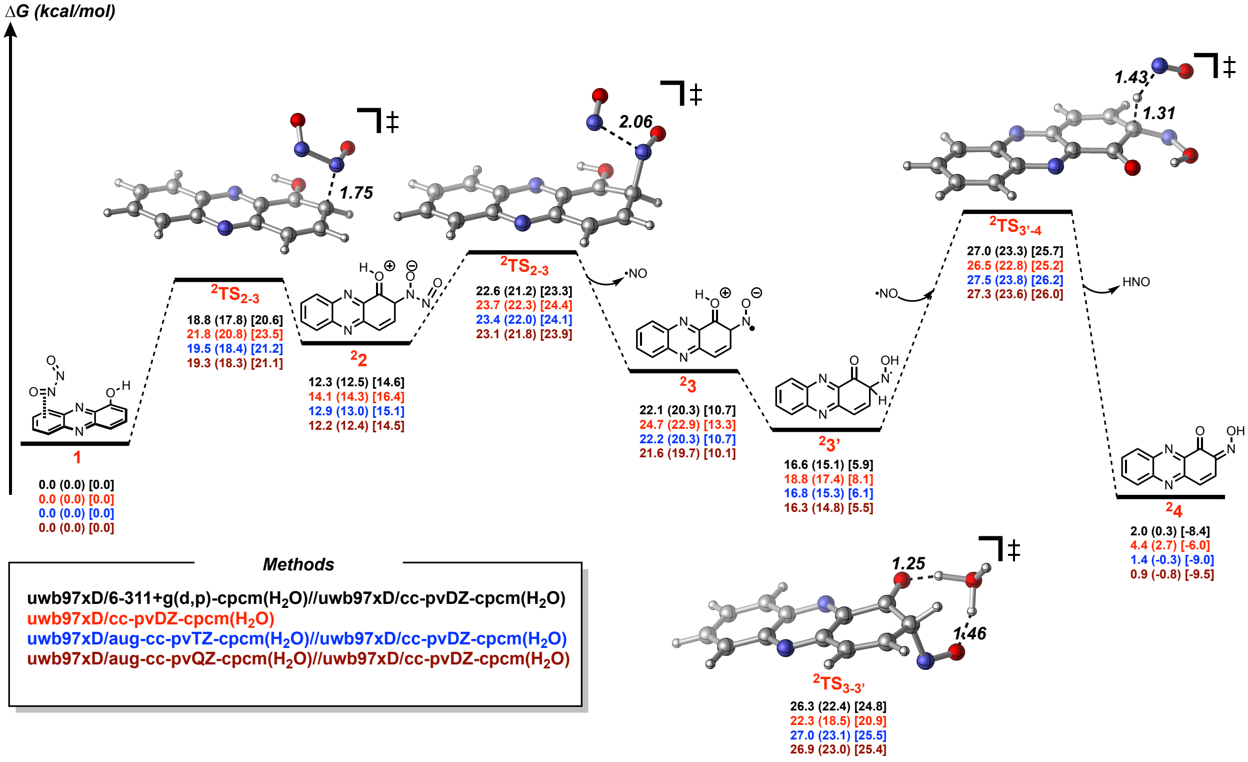


**Figure S5**. Energetic pathway of 1,2-addition of Nitric oxide to 1-Hydroxyphenazine calculated at different computational methods.

- *$^{2}TS_{3\text{-}3'}$ represents the tautomerization between intermediates $^{2}3$ and $^{2}3'$*

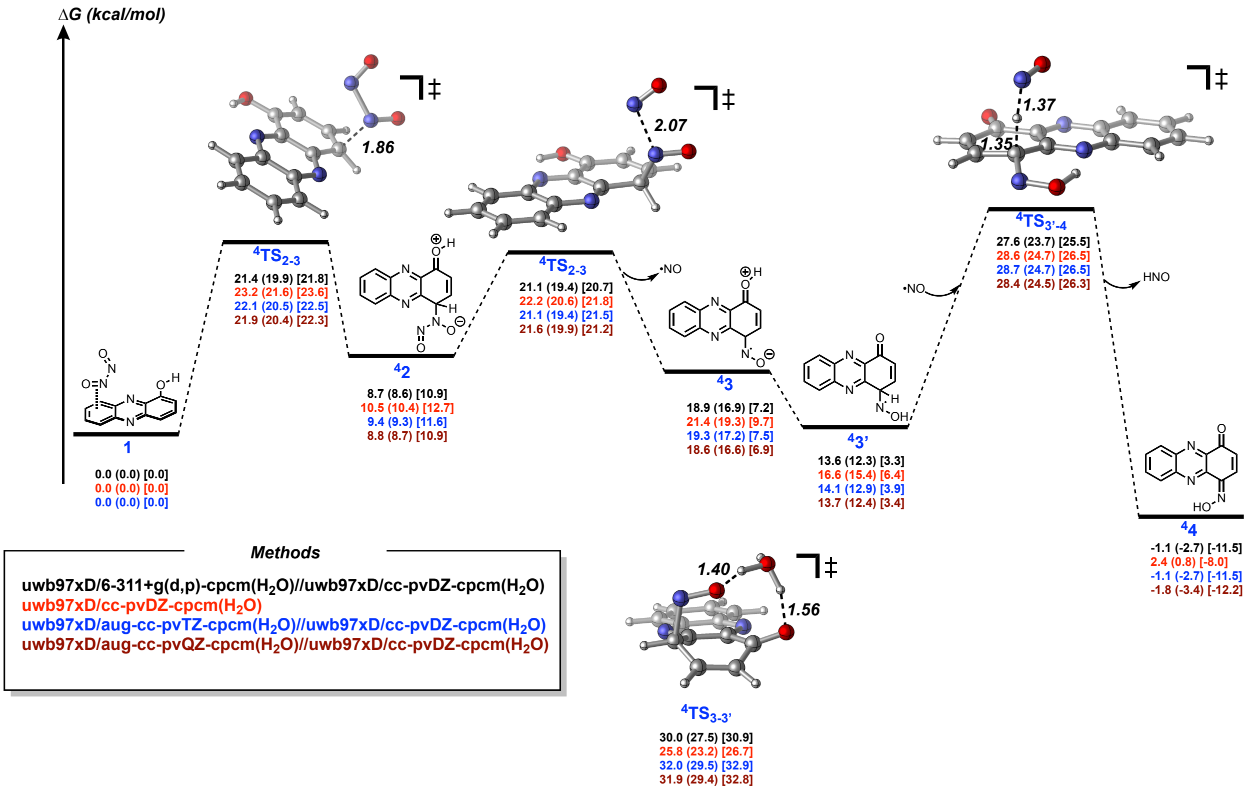


**Figure S6**. Energetic pathway of 1,4-addition of Nitric oxide to 1-Hydroxyphenazine calculated at different computational methods.

- *$^{4}TS_{3\text{-}3'}$ represents the tautomerization between intermediates $^{4}3$ and $^{4}3'$*

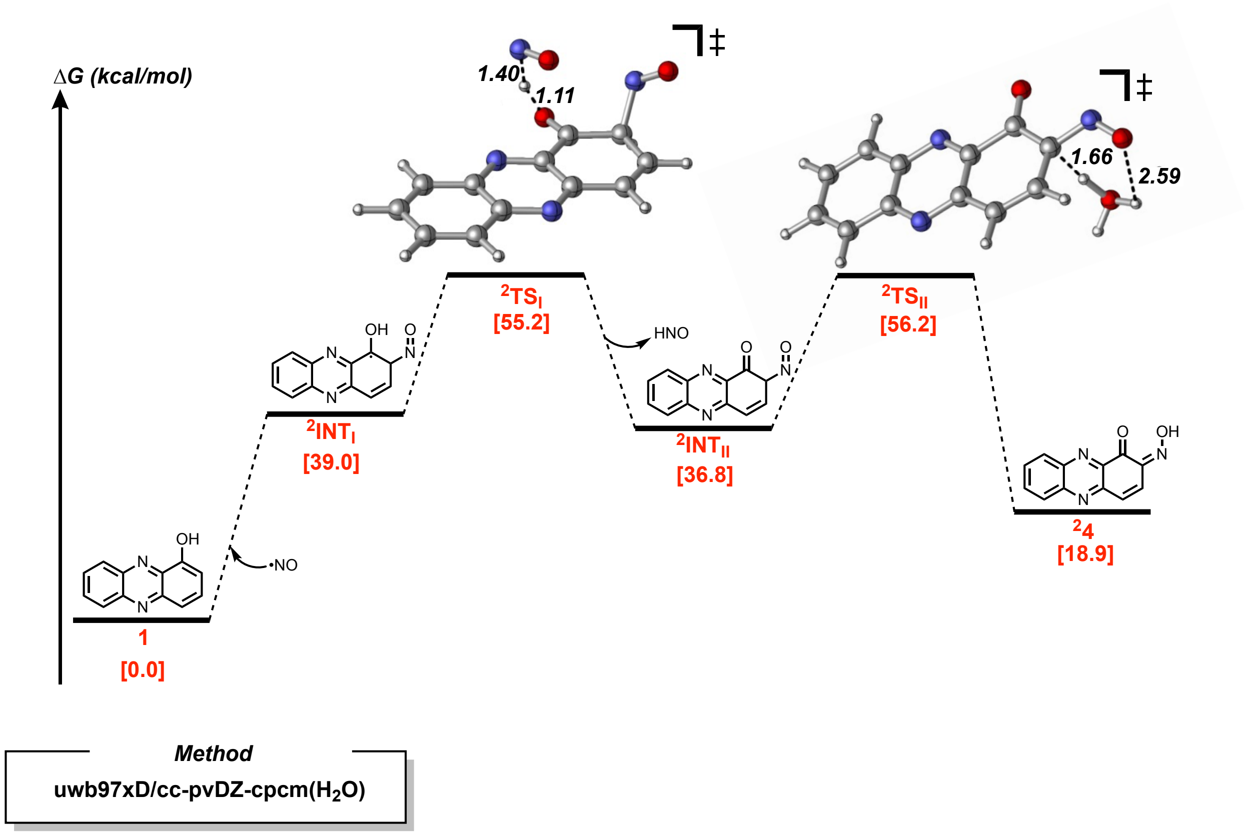


**Figure S7**. Alternative mechanism of 1,2-addition of Nitric oxide to 1-HydroxyPhenazine.

- ❑ *Addition of Nitric Oxide to the Phenazine ring was determined to be a completely uphill process.*

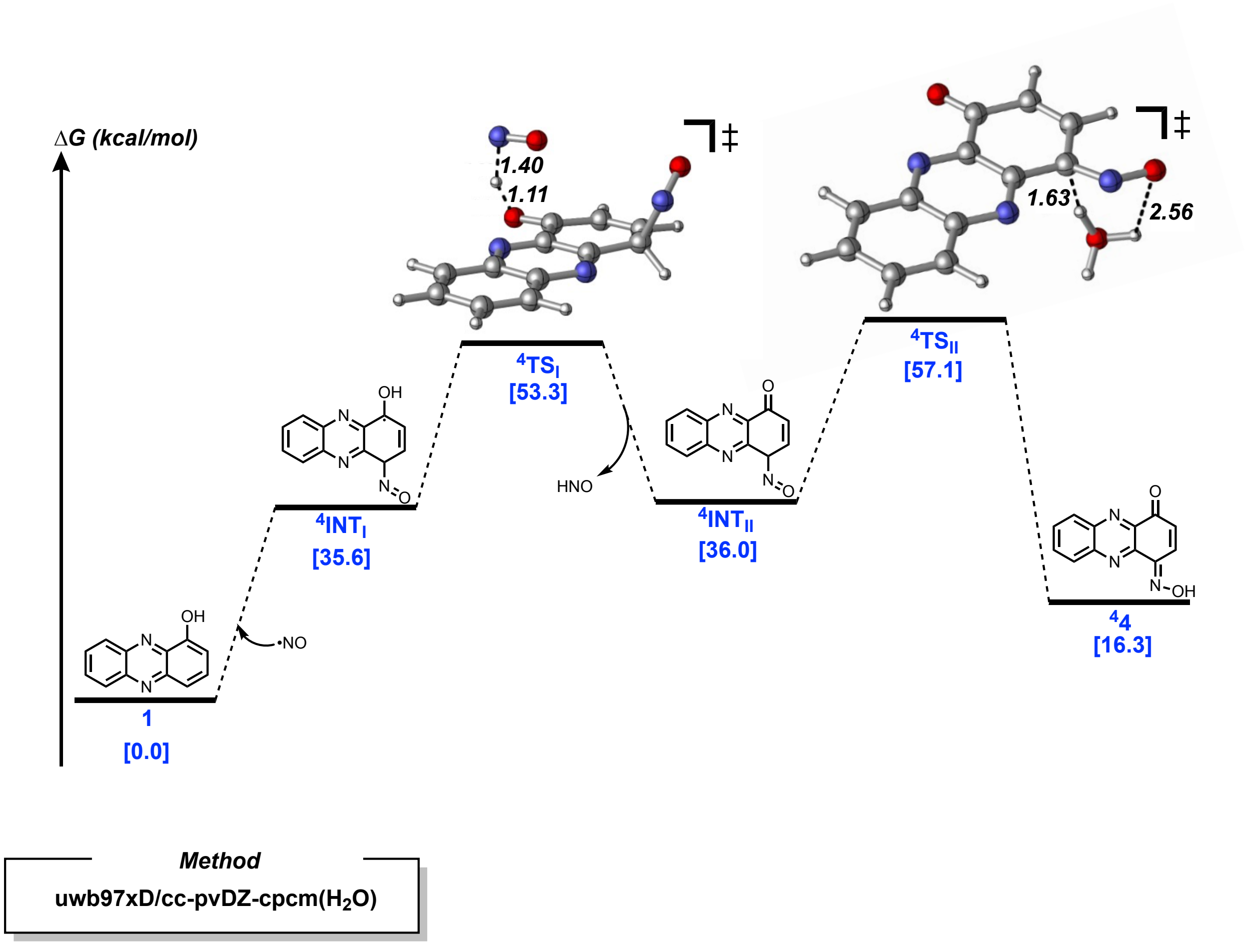


**Figure S8**. Alternative mechanism of 1,4-addition of Nitric oxide to 1-Hydroxyphenazine.

- *Addition of Nitric Oxide to the Phenazine ring was determined to be a completely uphill process.*

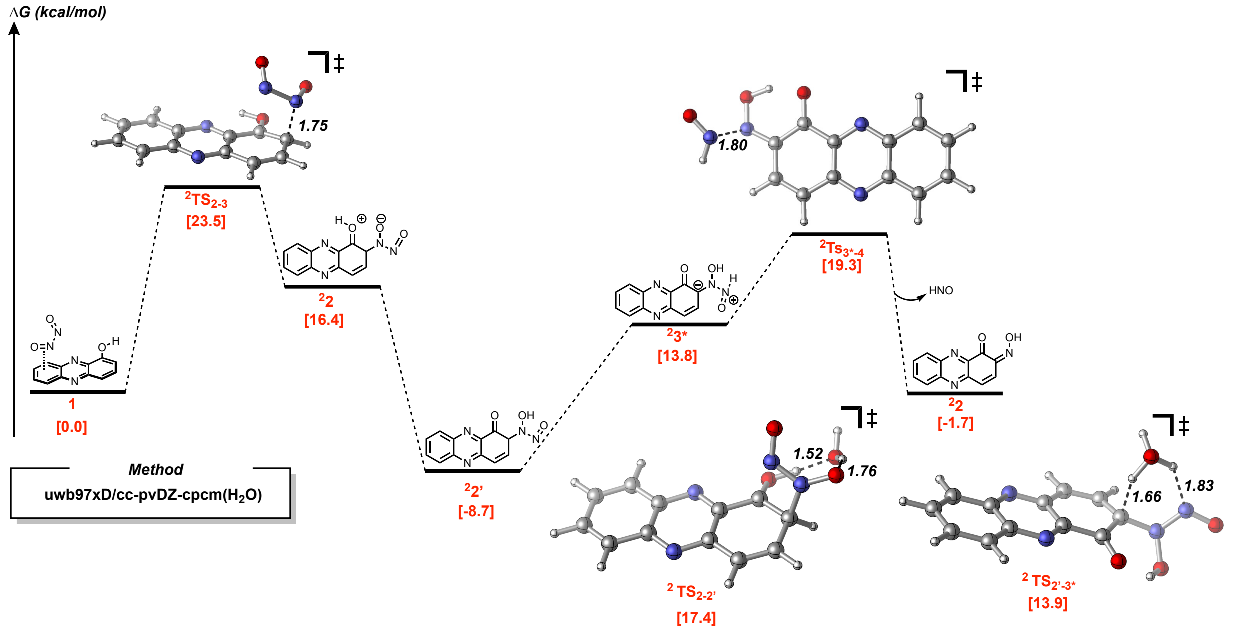

**Figure S9**. Alternative mechanism of 1,2-addition of Nitric oxide to 1-HydroxyPhenazine.

- ❑ *$^{2}TS_{2\text{-}2'}$ represents the tautomerization between intermediates $^{2}2$ and $^{2}2'$*
- ❑ *$^{2}TS_{2'\text{-}3*}$ represents the tautomerization between intermediates $^{2}2'$ and $^{2}3*$*

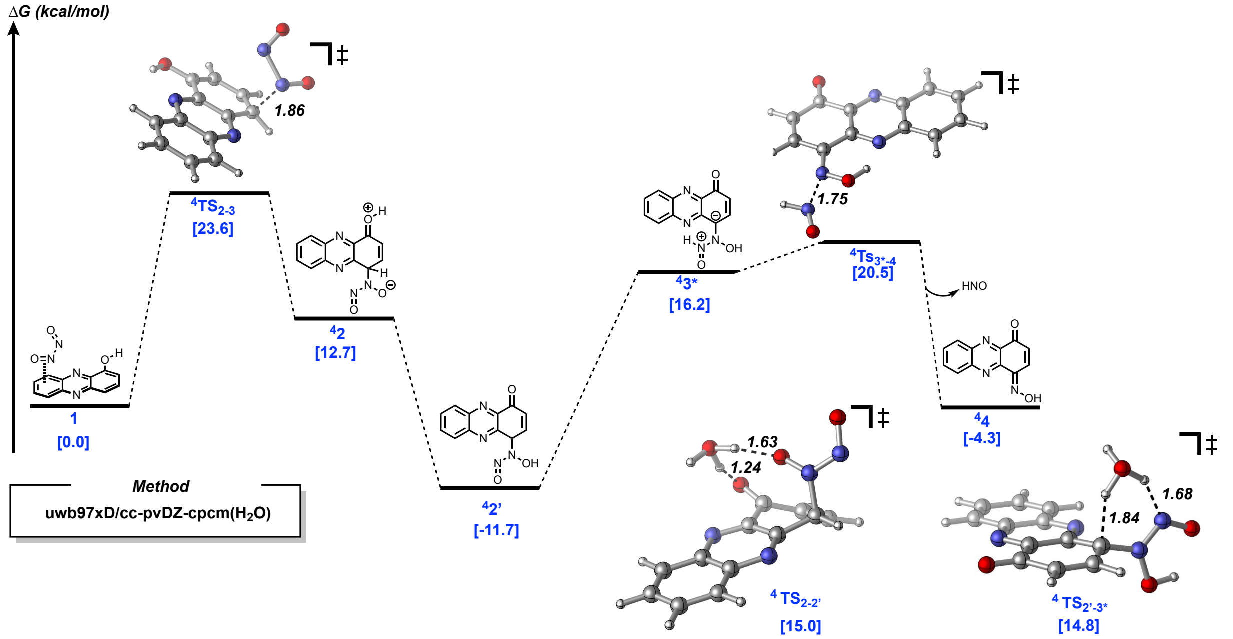

**Figure S10**. Alternative mechanism of 1,4-addition of Nitric oxide to 1-HydroxyPhenazine.

- ❑ *$^{4}TS_{2\text{-}2'}$ represents the tautomerization between intermediates $^{4}2$ and $^{4}2'$*
- ❑ *$^{4}TS_{2'\text{-}3*}$ represents the tautomerization between intermediates $^{4}2'$ and $^{4}3*$*

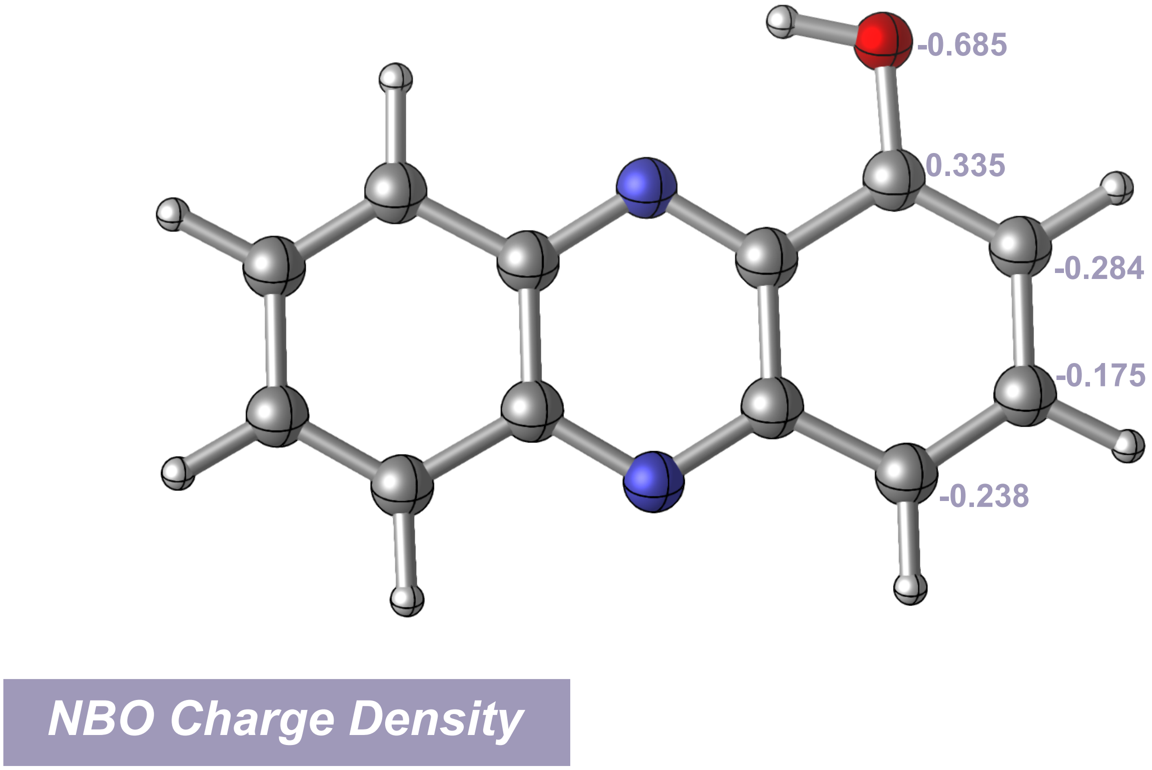


**Figure S11**. NBO charge density of 1-Hydroxyphenazine at the uwb97xD/6-311+g(d,p)-cpcm($H_2O$)//uwb97xD//cc-pvDZ-cpcm($H_2O$) level of theory.

- *The 2 and 4 positions represent the most nucleophilic sites of the ring.*

**Table S1.** Cartesian coordinates (xyz format) and energies of all the structures involved in each reaction mechanism studied calculated at the uwb97xD//cc-pvDZ-cpcm($H_2O$) level of theory.

**1**

E(scf) = -906.381162860 a.u.

| | | | | | | | |
|---|---|---|---|---|---|---|---|
| C | -4.095080 | -0.375167 | -0.467441 | C | 1.916123 | 1.457707 | 0.108544 |
| C | -2.995123 | -0.788045 | -1.162784 | C | 0.636314 | 0.887111 | -0.230215 |
| C | -2.778965 | 1.176761 | 0.862851 | H | 1.651383 | -1.250665 | -2.730760 |
| C | -3.986394 | 0.617811 | 0.556780 | H | 3.834818 | -0.222824 | -2.127660 |
| H | -5.073578 | -0.803589 | -0.689306 | H | 4.004471 | 1.483868 | -0.320833 |
| H | -3.056063 | -1.543967 | -1.946593 | N | -0.420761 | 1.314119 | 0.462348 |
| H | -2.671872 | 1.934015 | 1.640412 | N | -0.642576 | -0.649873 | -1.567060 |
| H | -4.883582 | 0.926231 | 1.095273 | O | 1.956516 | 2.373396 | 1.086983 |
| C | -1.709433 | -0.229056 | -0.874135 | H | 1.036414 | 2.472539 | 1.396156 |
| C | -1.604951 | 0.769720 | 0.156443 | N | 2.231258 | -1.320397 | 1.506419 |
| C | 0.541206 | -0.102392 | -1.261422 | N | 0.404999 | -1.216261 | 2.046500 |
| C | 1.729644 | -0.496780 | -1.947797 | O | -0.080574 | -2.012772 | 1.368440 |
| C | 2.925043 | 0.077472 | -1.605373 | O | 2.182325 | -2.133738 | 0.691748 |
| C | 3.035191 | 1.054220 | -0.574035 | | | | |

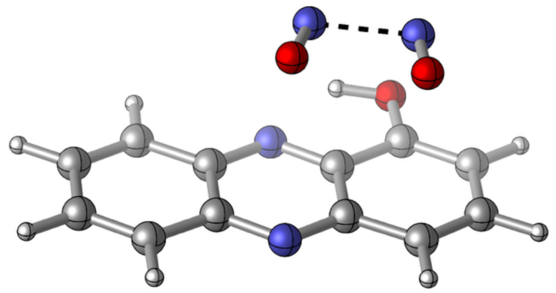

Zero-point correction= 0.190614 (Hartree/Particle)
Thermal correction to Energy= 0.206516
Thermal correction to Enthalpy= 0.207460
Thermal correction to Gibbs Free Energy= 0.145511
Sum of electronic and zero-point Energies= -906.190549
Sum of electronic and thermal Energies= -906.174647
Sum of electronic and thermal Enthalpies= -906.173703
Sum of electronic and thermal Free Energies= -906.235652

**$^{2}TS_{2\text{-}3}$**
E(scf) = -906.346379561 a.u.
$\nu_{min}$ = -546.0 $cm^{-1}$

| | | | | | | | |
|---|---|---|---|---|---|---|---|
| C | -0.585584 | -0.831972 | 0.010453 | C | -6.721352 | -1.074683 | -1.424717 |
| C | -1.548264 | 0.140499 | 0.114139 | C | -5.367858 | -0.775053 | -1.018337 |
| C | -2.191944 | -2.425595 | -0.868689 | H | -5.845067 | 2.523513 | -0.116341 |
| C | -0.909938 | -2.124509 | -0.484629 | H | -8.173115 | 1.957316 | -0.722453 |
| H | 0.441135 | -0.616375 | 0.309375 | N | -4.475079 | -1.738633 | -1.146751 |
| H | -1.320996 | 1.138498 | 0.490773 | N | -3.823189 | 0.840173 | -0.163116 |
| H | -2.463428 | -3.410237 | -1.251180 | O | -6.993657 | -2.264637 | -1.926634 |
| H | -0.127389 | -2.880950 | -0.557930 | H | -6.169395 | -2.789518 | -1.855668 |
| C | -2.886434 | -0.137806 | -0.273032 | H | -8.642025 | -0.212330 | -1.876600 |
| C | -3.209639 | -1.439017 | -0.770501 | N | -8.466087 | -1.025109 | 0.131351 |
| C | -5.052780 | 0.533288 | -0.527951 | O | -9.061411 | -2.043736 | -0.142816 |
| C | -6.117352 | 1.521425 | -0.449117 | N | -7.241712 | -1.238322 | 1.107847 |
| C | -7.383515 | 1.208419 | -0.783992 | O | -7.313937 | -2.315499 | 1.582163 |
| C | -7.778550 | -0.145242 | -1.211217 | | | | |

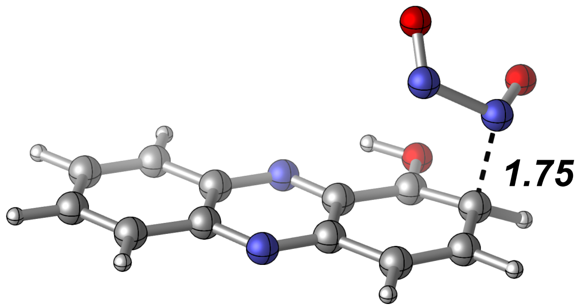


Zero-point correction= 0.190319 (Hartree/Particle)
Thermal correction to Energy= 0.204846
Thermal correction to Enthalpy= 0.205790
Thermal correction to Gibbs Free Energy= 0.148215
Sum of electronic and zero-point Energies= -906.156060
Sum of electronic and thermal Energies= -906.141534
Sum of electronic and thermal Enthalpies= -906.140589
Sum of electronic and thermal Free Energies= -906.198165

$^{2}$**2**

E(scf) = -906.358671158 a.u.

| | | | |
|---|---|---|---|
| C | 4.848731 | 0.252082 | 0.553838 |
| C | 3.823889 | 1.162580 | 0.395969 |
| C | 3.339720 | -1.591419 | 0.103220 |
| C | 4.602748 | -1.132751 | 0.405761 |
| H | 5.854778 | 0.599174 | 0.793519 |
| H | 3.990004 | 2.235496 | 0.504010 |
| H | 3.129346 | -2.655447 | -0.014654 |
| H | 5.422215 | -1.841824 | 0.533089 |
| C | 2.519911 | 0.720619 | 0.086797 |
| C | 2.269776 | -0.676293 | -0.061647 |
| C | 0.329484 | 1.198098 | -0.349641 |
| C | -0.785377 | 2.127075 | -0.542276 |
| C | -2.028331 | 1.702244 | -0.800509 |
| C | -2.391869 | 0.248746 | -0.932180 |
| C | -1.222360 | -0.666470 | -0.813330 |
| C | 0.071187 | -0.216236 | -0.490767 |
| H | -0.558185 | 3.191209 | -0.465362 |
| H | -2.845948 | 2.412168 | -0.933034 |
| N | 1.026801 | -1.141156 | -0.355925 |
| N | 1.520324 | 1.653649 | -0.069717 |
| O | -1.453107 | -1.964803 | -0.988080 |
| H | -0.597171 | -2.409769 | -0.820114 |
| H | -2.936724 | 0.065509 | -1.870898 |
| N | -3.386684 | -0.138996 | 0.116407 |
| O | -4.376697 | -0.824081 | -0.137355 |
| N | -2.995312 | 0.177683 | 1.445154 |
| O | -3.780105 | -0.239171 | 2.249260 |

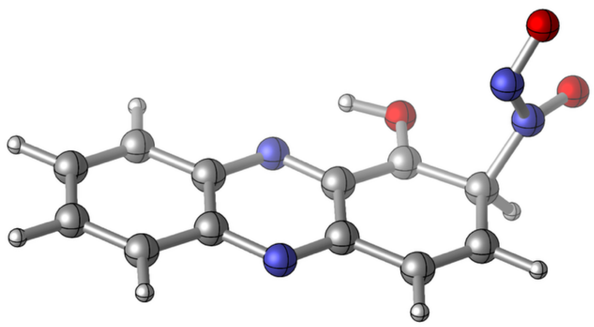

Zero-point correction= 0.192243 (Hartree/Particle)
Thermal correction to Energy= 0.206799
Thermal correction to Enthalpy= 0.207743
Thermal correction to Gibbs Free Energy= 0.149144
Sum of electronic and zero-point Energies= -906.166428
Sum of electronic and thermal Energies= -906.151872
Sum of electronic and thermal Enthalpies= -906.150928
Sum of electronic and thermal Free Energies= -906.209527

**$^{2}TS_{2\text{-}3}$**

E(scf) = -906.343441288 a.u.

$\nu_{min}$ = -259.0 $cm^{-1}$

| | | | | | | | |
|---|---|---|---|---|---|---|---|
| C | 0.080045 | 1.052552 | 0.002202 | C | -6.026591 | 0.822488 | -1.529219 |
| C | -0.900422 | 2.015746 | 0.108970 | C | -4.705211 | 1.105433 | -1.103139 |
| C | -1.507056 | -0.523608 | -0.934610 | H | -5.182513 | 4.403828 | -0.166124 |
| C | -0.227545 | -0.224965 | -0.523403 | H | -7.499693 | 3.867271 | -0.802641 |
| H | 1.099196 | 1.272990 | 0.322944 | N | -3.794879 | 0.141916 | -1.237349 |
| H | -0.687167 | 3.007890 | 0.509796 | N | -3.176645 | 2.718155 | -0.189335 |
| H | -1.764300 | -1.502411 | -1.342249 | O | -6.304629 | -0.374992 | -2.041502 |
| H | 0.558371 | -0.977763 | -0.602465 | H | -5.470778 | -0.884885 | -1.984520 |
| C | -2.222789 | 1.736615 | -0.303725 | H | -7.880628 | 1.748927 | -2.104759 |
| C | -2.533558 | 0.449150 | -0.833184 | N | -7.905279 | 1.065552 | -0.087844 |
| C | -4.387524 | 2.411752 | -0.577779 | O | -8.553843 | 0.098483 | -0.390388 |
| C | -5.458478 | 3.405308 | -0.508474 | N | -6.544196 | 0.348410 | 1.287748 |
| C | -6.717455 | 3.109212 | -0.863871 | O | -6.951125 | -0.707290 | 1.431226 |
| C | -7.142742 | 1.743029 | -1.290156 | | | | |

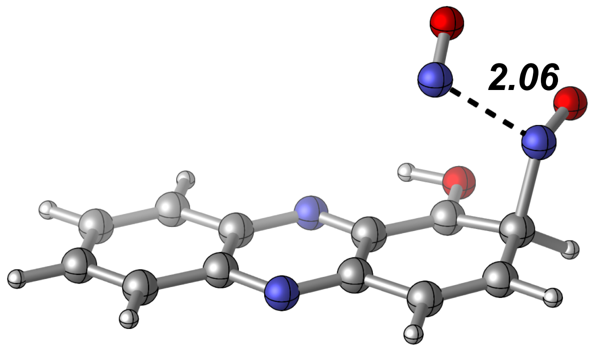


Zero-point correction= 0.189535 (Hartree/Particle)
Thermal correction to Energy= 0.204358
Thermal correction to Enthalpy= 0.205302
Thermal correction to Gibbs Free Energy= 0.146643
Sum of electronic and zero-point Energies= -906.153906
Sum of electronic and thermal Energies= -906.139084
Sum of electronic and thermal Enthalpies= -906.138139
Sum of electronic and thermal Free Energies= -906.196798

**$^{2}$3**

E(scf) = -776.483897779 a.u.

| | | | | | | | |
|---|---|---|---|---|---|---|---|
| C | 4.396090 | 0.266549 | 0.335634 | C | -2.931280 | 0.243789 | -0.388666 |
| C | 3.358863 | 1.174143 | 0.288629 | C | -1.775791 | -0.653908 | -0.510334 |
| C | 2.861229 | -1.574615 | -0.029403 | C | -0.452761 | -0.207277 | -0.292838 |
| C | 4.143425 | -1.116390 | 0.174649 | H | -1.083557 | 3.197570 | -0.227471 |
| H | 5.417903 | 0.613313 | 0.496290 | H | -3.400836 | 2.401021 | -0.490592 |
| H | 3.529659 | 2.245210 | 0.407595 | N | 0.514822 | -1.126636 | -0.273237 |
| H | 2.645281 | -2.636685 | -0.154992 | N | 1.022632 | 1.662606 | 0.030020 |
| H | 4.973686 | -1.823350 | 0.213021 | O | -2.010447 | -1.952419 | -0.692173 |
| C | 2.033190 | 0.732569 | 0.082077 | H | -1.139577 | -2.391622 | -0.609206 |
| C | 1.777065 | -0.661942 | -0.078308 | H | -3.755206 | -0.026992 | -1.062959 |
| C | -0.189204 | 1.206366 | -0.152037 | N | -3.466763 | -0.106646 | 1.074154 |
| C | -1.317115 | 2.132225 | -0.256796 | O | -4.195984 | -1.046002 | 1.070072 |
| C | -2.575503 | 1.692429 | -0.407846 | | | | |

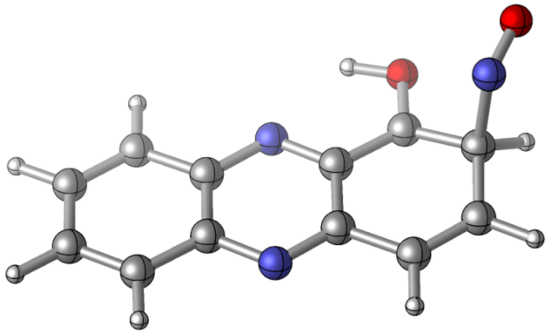

Zero-point correction= 0.182890 (Hartree/Particle)
Thermal correction to Energy= 0.195597
Thermal correction to Enthalpy= 0.196541
Thermal correction to Gibbs Free Energy= 0.142617
Sum of electronic and zero-point Energies= -776.301008
Sum of electronic and thermal Energies= -776.288301
Sum of electronic and thermal Enthalpies= -776.287357
Sum of electronic and thermal Free Energies= -776.341281

**$^2$TS$_{3\text{-}3'}$**

E(scf) = -852.893830909 a.u.

$\nu_{min}$ = -1011.4 $cm^{-1}$

| | | | | | | | |
|---|---|---|---|---|---|---|---|
| C | 0.042159 | 0.085430 | -0.002905 | C | -7.225060 | 0.973683 | 0.540759 |
| C | -0.859420 | 1.116641 | 0.101852 | C | -6.258431 | -0.119824 | 0.755090 |
| C | -1.718331 | -1.567742 | 0.212055 | C | -4.822454 | 0.178857 | 0.558667 |
| C | -0.388727 | -1.266889 | 0.052382 | H | -4.993393 | 3.638977 | 0.702695 |
| H | 1.103799 | 0.302769 | -0.129764 | H | -7.421925 | 3.157129 | 0.792887 |
| H | -0.547215 | 2.160929 | 0.062478 | N | -3.983993 | -0.831121 | 0.479000 |
| H | -2.076519 | -2.596954 | 0.258188 | N | -3.122986 | 1.865364 | 0.371878 |
| H | 0.347044 | -2.067657 | -0.032707 | O | -6.606377 | -1.307467 | 0.982282 |
| C | -2.240436 | 0.835775 | 0.265470 | N | -7.456225 | 0.744619 | -0.997577 |
| C | -2.672570 | -0.521194 | 0.321439 | O | -8.334116 | -0.151131 | -1.221364 |
| C | -4.394661 | 1.549518 | 0.510776 | H | -7.840519 | -1.512477 | 0.967076 |
| C | -5.380655 | 2.620830 | 0.646893 | O | -8.955672 | -1.632599 | 0.664917 |
| C | -6.691413 | 2.352162 | 0.702329 | | | | |

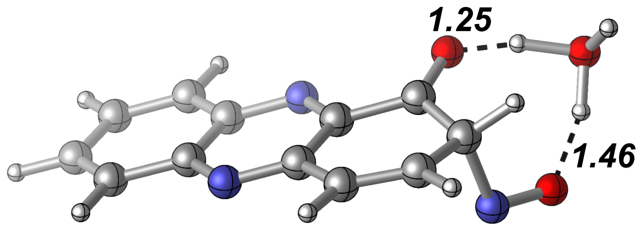


Zero-point correction= 0.203767 (Hartree/Particle)
Thermal correction to Energy= 0.217685
Thermal correction to Enthalpy= 0.218629
Thermal correction to Gibbs Free Energy= 0.162354
Sum of electronic and zero-point Energies= -852.690064
Sum of electronic and thermal Energies= -852.676146
Sum of electronic and thermal Enthalpies= -852.675202
Sum of electronic and thermal Free Energies= -852.731477

**[2]3’**

E(scf) = -776.493289570 a.u.

| | | | | | | | |
|---|---|---|---|---|---|---|---|
| C | -4.498926 | 0.203047 | -0.174986 | C | 2.893601 | 0.336320 | 0.359214 |
| C | -3.487497 | 1.131568 | -0.159786 | C | 1.774383 | -0.697636 | 0.274957 |
| C | -2.921929 | -1.628788 | 0.017638 | C | 0.363741 | -0.221327 | 0.157894 |
| C | -4.216363 | -1.187065 | -0.086020 | H | 0.916732 | 3.195333 | -0.029685 |
| H | -5.536094 | 0.531667 | -0.256388 | H | 3.274368 | 2.499169 | 0.174876 |
| H | -3.686031 | 2.201773 | -0.227085 | N | -0.579730 | -1.136022 | 0.140324 |
| H | -2.675817 | -2.688939 | 0.087658 | N | -1.144480 | 1.631340 | -0.040048 |
| H | -5.039352 | -1.902656 | -0.100795 | O | 2.046393 | -1.886722 | 0.317790 |
| C | -2.138025 | 0.704876 | -0.053429 | H | 3.303569 | 0.217293 | 1.385782 |
| C | -1.855121 | -0.690359 | 0.036134 | N | 3.995561 | 0.082709 | -0.588532 |
| C | 0.090490 | 1.180690 | 0.066038 | O | 4.429118 | -1.187620 | -0.534075 |
| C | 1.192489 | 2.144959 | 0.072080 | H | 3.749696 | -1.744748 | -0.081440 |
| C | 2.468055 | 1.763657 | 0.189457 | | | | |

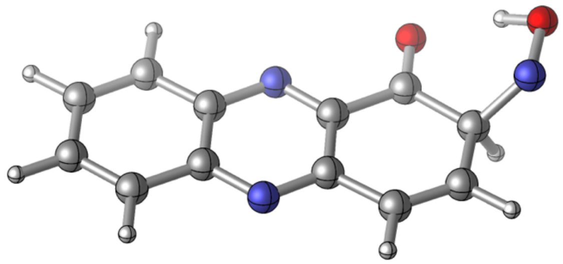

Zero-point correction= 0.183897 (Hartree/Particle)
Thermal correction to Energy= 0.196224
Thermal correction to Enthalpy= 0.197168
Thermal correction to Gibbs Free Energy= 0.143756
Sum of electronic and zero-point Energies= -776.309392
Sum of electronic and thermal Energies= -776.297066
Sum of electronic and thermal Enthalpies= -776.296122
Sum of electronic and thermal Free Energies= -776.349534

**$^{2}$TS$_{3'-4}$**
E(scf) = -906.3389236132 a.u.
$\nu_{min}$ = -1651.8 cm$^{-1}$

| | | | | | | | |
|---|---|---|---|---|---|---|---|
| C | 0.305322 | -0.819161 | -0.228938 | C | 7.665586 | -0.627433 | -0.111248 |
| C | 1.302171 | 0.104616 | -0.040007 | C | 6.604256 | -1.609608 | -0.505975 |
| C | 1.915174 | -2.584471 | -0.646269 | C | 5.186325 | -1.157966 | -0.371781 |
| C | 0.611894 | -2.174139 | -0.534500 | H | 5.688265 | 2.209688 | 0.282064 |
| H | -0.739226 | -0.515495 | -0.144899 | H | 8.069021 | 1.505562 | 0.182641 |
| H | 1.085606 | 1.147362 | 0.194762 | N | 4.252836 | -2.063923 | -0.568589 |
| H | 2.179264 | -3.616540 | -0.879743 | N | 3.640326 | 0.630412 | 0.040145 |
| H | -0.200818 | -2.886865 | -0.679981 | O | 6.881482 | -2.744710 | -0.877708 |
| C | 2.662803 | -0.290091 | -0.148120 | H | 7.779738 | -1.023398 | 1.135374 |
| C | 2.969592 | -1.649675 | -0.455288 | N | 8.956168 | -0.715662 | -0.623190 |
| C | 4.888948 | 0.209518 | -0.064167 | O | 9.322647 | -1.938473 | -1.045901 |
| C | 5.972104 | 1.170354 | 0.115946 | H | 8.532684 | -2.537022 | -1.014531 |
| C | 7.255650 | 0.788125 | 0.062633 | | | | |

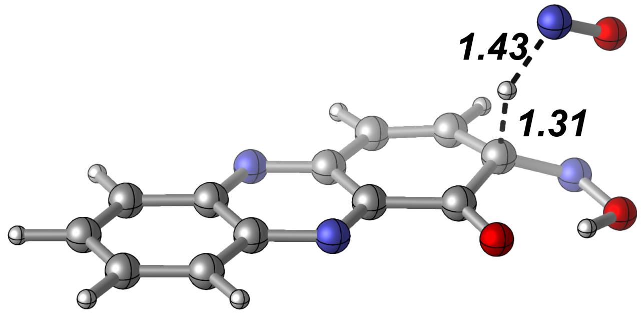


Zero-point correction= 0.185967 (Hartree/Particle)
Thermal correction to Energy= 0.200653
Thermal correction to Enthalpy= 0.201597
Thermal correction to Gibbs Free Energy= 0.143486
Sum of electronic and zero-point Energies= -906.152956
Sum of electronic and thermal Energies= -906.138271
Sum of electronic and thermal Enthalpies= -906.137327
Sum of electronic and thermal Free Energies= -906.195438

**$^{2}$4**

E(scf) = -775.932953859 a.u.

| | | | | | | | |
|---|---|---|---|---|---|---|---|
| C | -4.485908 | 0.189889 | 0.000117 | C | 2.482679 | 1.765458 | -0.000266 |
| C | -3.480996 | 1.123895 | 0.000006 | C | 2.852665 | 0.356165 | -0.000138 |
| C | -2.892235 | -1.638600 | 0.000247 | C | 1.803778 | -0.686257 | 0.000024 |
| C | -4.191494 | -1.202052 | 0.000237 | C | 0.391051 | -0.214626 | 0.000025 |
| H | -5.527871 | 0.513453 | 0.000113 | H | 0.919356 | 3.212889 | -0.000339 |
| H | -3.688300 | 2.194532 | -0.000089 | H | 3.301802 | 2.485557 | -0.000373 |
| H | -2.637479 | -2.699017 | 0.000338 | N | -0.550289 | -1.134581 | 0.000143 |
| H | -5.010888 | -1.921927 | 0.000322 | N | -1.138646 | 1.634237 | -0.000111 |
| C | -2.123568 | 0.703349 | 0.000009 | O | 2.087124 | -1.885276 | 0.000144 |
| C | -1.828935 | -0.693994 | 0.000134 | N | 4.134431 | 0.091429 | -0.000171 |
| C | 0.106763 | 1.189880 | -0.000108 | O | 4.517919 | -1.175419 | -0.000054 |
| C | 1.197010 | 2.158880 | -0.000247 | H | 3.699688 | -1.749429 | 0.000048 |

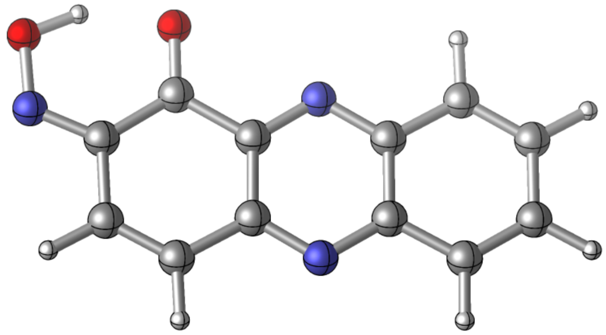

Zero-point correction= 0.174156 (Hartree/Particle)
Thermal correction to Energy= 0.185794
Thermal correction to Enthalpy= 0.186738
Thermal correction to Gibbs Free Energy= 0.135980
Sum of electronic and zero-point Energies= -775.758798
Sum of electronic and thermal Energies= -775.747160
Sum of electronic and thermal Enthalpies= -775.746216
Sum of electronic and thermal Free Energies= -775.796973

**$^{4}$TS$_{2\text{-}3}$**
E(scf) = -906.344243727 a.u.
$\nu_{min}$ = -603.0 cm$^{-1}$

| | | | | | | | |
|---|---|---|---|---|---|---|---|
| C | -0.970555 | -0.043796 | 0.081656 | C | 4.309941 | 3.369396 | -0.508450 |
| C | 0.256584 | -0.412204 | -0.406809 | C | 3.259191 | 2.376140 | -0.438285 |
| C | -0.183914 | 2.203286 | 0.562505 | H | 6.751114 | 1.504209 | -1.954227 |
| C | -1.191644 | 1.274121 | 0.571018 | H | 6.303257 | 3.807946 | -1.147672 |
| H | -1.789534 | -0.764173 | 0.098908 | N | 2.089340 | 2.769908 | 0.040907 |
| H | 0.449355 | -1.416987 | -0.784606 | N | 2.535352 | 0.140059 | -0.901934 |
| H | -0.332372 | 3.219204 | 0.930505 | O | 4.035279 | 4.595571 | -0.057897 |
| H | -2.176686 | 1.542592 | 0.955384 | H | 3.108367 | 4.565211 | 0.250443 |
| C | 1.325133 | 0.525766 | -0.428208 | H | 4.890089 | -0.175540 | -2.035734 |
| C | 1.099268 | 1.851259 | 0.059629 | O | 6.528768 | -0.955736 | -0.135391 |
| C | 3.494704 | 1.044453 | -0.903575 | N | 5.551793 | -0.290306 | 0.071620 |
| C | 4.844220 | 0.671436 | -1.348417 | N | 5.768596 | 0.811912 | 1.277488 |
| C | 5.781203 | 1.736955 | -1.513127 | O | 6.743870 | 0.543730 | 1.860960 |
| C | 5.534051 | 3.040222 | -1.061600 | | | | |

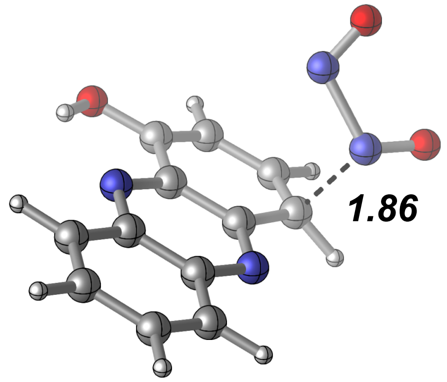


Zero-point correction= 0.189159 (Hartree/Particle)
Thermal correction to Energy= 0.204039
Thermal correction to Enthalpy= 0.204983
Thermal correction to Gibbs Free Energy= 0.146159
Sum of electronic and zero-point Energies= -906.155085
Sum of electronic and thermal Energies= -906.140205
Sum of electronic and thermal Enthalpies= -906.139261
Sum of electronic and thermal Free Energies= -906.198084

**$^{4}$2**

E(scf) = -906.364453747 a.u.

| | | | |
|---|---|---|---|
| C | -4.040079 | -1.546030 | -0.138133 |
| C | -2.734550 | -1.781168 | -0.508746 |
| C | -3.501418 | 0.730145 | 0.508707 |
| C | -4.423226 | -0.282460 | 0.374040 |
| H | -4.786326 | -2.335362 | -0.237675 |
| H | -2.414266 | -2.745945 | -0.904420 |
| H | -3.777828 | 1.710144 | 0.899956 |
| H | -5.461516 | -0.113374 | 0.663509 |
| C | -1.767250 | -0.758772 | -0.379055 |
| C | -2.151825 | 0.513528 | 0.133082 |
| C | 0.385563 | -0.042952 | -0.605026 |
| C | 1.832227 | -0.310121 | -0.976131 |
| C | 2.733730 | 0.885456 | -0.875946 |
| C | 2.307331 | 2.090718 | -0.396037 |
| C | 0.969000 | 2.295421 | 0.005360 |
| C | 0.001409 | 1.244667 | -0.103242 |
| H | 3.768903 | 0.743932 | -1.186840 |
| H | 3.000311 | 2.930391 | -0.324051 |
| N | -1.244429 | 1.517923 | 0.261536 |
| N | -0.471232 | -1.013074 | -0.747935 |
| O | 0.566710 | 3.478471 | 0.478190 |
| H | -0.386485 | 3.375471 | 0.670009 |
| H | 1.871510 | -0.762132 | -1.976633 |
| O | 2.941671 | -2.384438 | -0.534687 |
| N | 2.375219 | -1.385341 | -0.095329 |
| N | 2.263628 | -1.111108 | 1.288307 |
| O | 2.748861 | -1.980597 | 1.958604 |

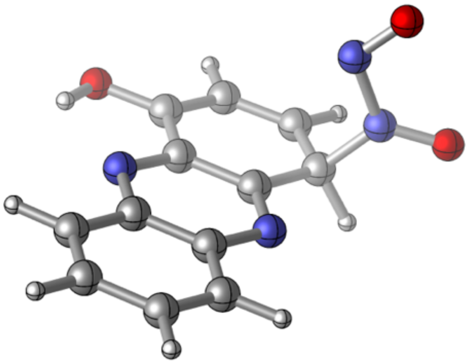

Zero-point correction= 0.191864 (Hartree/Particle)
Thermal correction to Energy= 0.206415
Thermal correction to Enthalpy= 0.207359
Thermal correction to Gibbs Free Energy= 0.148962
Sum of electronic and zero-point Energies= -906.172589
Sum of electronic and thermal Energies= -906.158039
Sum of electronic and thermal Enthalpies= -906.157095
Sum of electronic and thermal Free Energies= -906.215491

**$^{4}TS_{2-3}$**

E(scf) = -906.345764023 a.u.

$\nu_{min}$ = -274.9 $cm^{-1}$

| | | | | | | | |
|---|---|---|---|---|---|---|---|
| C | -1.115227 | -0.023313 | -0.002229 | C | 4.064538 | 3.579676 | -0.183700 |
| C | 0.158393 | -0.323038 | -0.433595 | C | 3.046562 | 2.575340 | -0.224341 |
| C | -0.440844 | 2.234790 | 0.574428 | H | 6.728024 | 1.862189 | -1.450852 |
| C | -1.414172 | 1.264386 | 0.505067 | H | 6.111579 | 4.103433 | -0.604564 |
| H | -1.900126 | -0.779406 | -0.049749 | N | 1.834281 | 2.914268 | 0.200643 |
| H | 0.414108 | -1.307919 | -0.827129 | N | 2.440021 | 0.331807 | -0.802132 |
| H | -0.652057 | 3.232774 | 0.961201 | O | 3.741772 | 4.789170 | 0.289270 |
| H | -2.427526 | 1.486339 | 0.843417 | H | 2.796256 | 4.730558 | 0.530543 |
| C | 1.177457 | 0.654552 | -0.372313 | H | 4.742754 | 0.414708 | -2.121313 |
| C | 0.877150 | 1.950835 | 0.135369 | O | 6.321268 | -0.668789 | -0.503395 |
| C | 3.349707 | 1.263329 | -0.725200 | N | 5.271264 | -0.165921 | -0.182609 |
| C | 4.755545 | 0.920046 | -1.142703 | N | 5.545136 | 0.687709 | 1.681271 |
| C | 5.709480 | 2.067570 | -1.120430 | O | 6.537945 | 0.238000 | 2.014883 |
| C | 5.371430 | 3.302379 | -0.640524 | | | | |

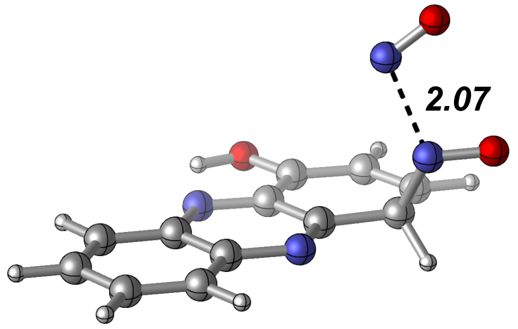


Zero-point correction= 0.188809 (Hartree/Particle)
Thermal correction to Energy= 0.203860
Thermal correction to Enthalpy= 0.204805
Thermal correction to Gibbs Free Energy= 0.144846
Sum of electronic and zero-point Energies= -906.156955
Sum of electronic and thermal Energies= -906.141904
Sum of electronic and thermal Enthalpies= -906.140959
Sum of electronic and thermal Free Energies= -906.200918

**[4]3**

E(scf) = -776.489168375 a.u.

| | | | | | | | |
|---|---|---|---|---|---|---|---|
| C | -4.046293 | -0.759610 | -0.128235 | C | 3.057636 | 0.854566 | -0.556451 |
| C | -2.859102 | -1.406026 | -0.385073 | C | 1.864024 | 1.459090 | -0.128694 |
| C | -2.878281 | 1.317145 | 0.331869 | C | 0.630926 | 0.716657 | -0.116167 |
| C | -4.053760 | 0.610747 | 0.234551 | H | 4.033660 | -0.942966 | -1.201518 |
| H | -4.990088 | -1.301739 | -0.201141 | H | 3.940451 | 1.483844 | -0.680613 |
| H | -2.827167 | -2.461020 | -0.660855 | N | -0.483274 | 1.382556 | 0.144105 |
| H | -2.863444 | 2.372971 | 0.605785 | N | -0.462142 | -1.374246 | -0.524936 |
| H | -5.004282 | 1.106934 | 0.436521 | O | 1.815110 | 2.758219 | 0.181995 |
| C | -1.633128 | -0.704670 | -0.289531 | H | 0.874959 | 2.948685 | 0.370289 |
| C | -1.641184 | 0.675331 | 0.066944 | H | 1.905297 | -2.275747 | -1.170253 |
| C | 0.642134 | -0.682285 | -0.430159 | O | 1.907997 | -1.353847 | 1.792601 |
| C | 1.958859 | -1.379616 | -0.542090 | N | 2.409371 | -1.927877 | 0.874772 |
| C | 3.107847 | -0.489492 | -0.849059 | | | | |

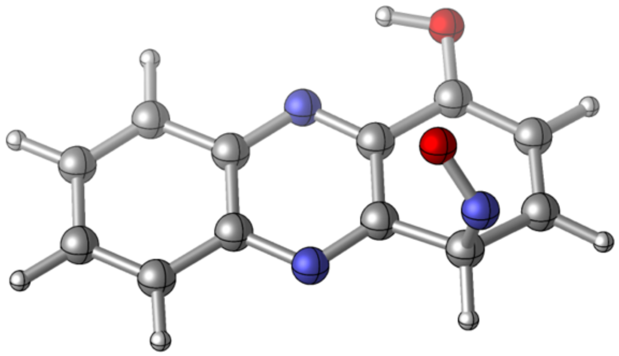

Zero-point correction= 0.182541 (Hartree/Particle)
Thermal correction to Energy= 0.195235
Thermal correction to Enthalpy= 0.196179
Thermal correction to Gibbs Free Energy= 0.142126
Sum of electronic and zero-point Energies= -776.306627
Sum of electronic and thermal Energies= -776.293933
Sum of electronic and thermal Enthalpies= -776.292989
Sum of electronic and thermal Free Energies= -776.347043

**[4]TS$_{3\text{-}3'}$**

E(scf) = -852.888360721 a.u.
$\nu_{min}$ = -198.1 $cm^{-1}$

| | | | | | | | |
|---|---|---|---|---|---|---|---|
| C | 0.041553 | -0.416108 | 0.012064 | C | 6.166986 | 0.904968 | -1.114118 |
| C | 1.144419 | -1.230202 | 0.093497 | C | 4.807733 | 0.386741 | -0.750455 |
| C | 1.410867 | 1.488587 | -0.600687 | H | 7.749214 | -2.121066 | -1.328426 |
| C | 0.174758 | 0.954754 | -0.335149 | H | 7.968618 | 0.223347 | -2.162247 |
| H | -0.950119 | -0.821419 | 0.218764 | N | 3.784773 | 1.207628 | -0.811547 |
| H | 1.063522 | -2.283568 | 0.364959 | N | 3.524368 | -1.516702 | -0.050735 |
| H | 1.540314 | 2.536704 | -0.873910 | O | 6.399729 | 2.127983 | -1.096371 |
| H | -0.715426 | 1.582826 | -0.390718 | O | 6.700773 | 0.163086 | 1.196145 |
| C | 2.436451 | -0.705399 | -0.172460 | N | 6.414496 | -1.080915 | 1.324966 |
| C | 2.568078 | 0.667795 | -0.531249 | H | 5.875866 | 2.511597 | 0.318398 |
| C | 4.684501 | -0.976441 | -0.343352 | O | 5.722551 | 2.382463 | 1.330597 |
| C | 5.961675 | -1.689029 | -0.091270 | H | 6.043755 | 1.365815 | 1.458737 |
| C | 6.997008 | -1.366607 | -1.091866 | H | 4.764517 | 2.425126 | 1.479897 |
| C | 7.122465 | -0.102389 | -1.555567 | H | 5.815343 | -2.756589 | 0.092369 |

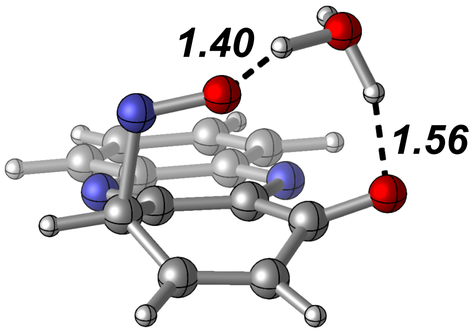


Zero-point correction= 0.206294 (Hartree/Particle)
Thermal correction to Energy= 0.219785
Thermal correction to Enthalpy= 0.220730
Thermal correction to Gibbs Free Energy= 0.166102
Sum of electronic and zero-point Energies= -852.682067
Sum of electronic and thermal Energies= -852.668575
Sum of electronic and thermal Enthalpies= -852.667631
Sum of electronic and thermal Free Energies= -852.722259

**$^{4}$3'**

E(scf) = -776.496791130 a.u.

| | | | | | | | |
|---|---|---|---|---|---|---|---|
| C | 3.939856 | 0.776187 | -0.073746 | C | -3.156268 | -1.178750 | -0.089342 |
| C | 2.713138 | 1.375776 | -0.217731 | C | -1.863720 | -1.874758 | 0.044248 |
| C | 2.926402 | -1.413646 | 0.181651 | C | -0.629593 | -1.012573 | -0.027751 |
| C | 4.048486 | -0.626736 | 0.126063 | H | -4.232712 | 0.619644 | -0.342531 |
| H | 4.847011 | 1.380675 | -0.113766 | H | -4.045718 | -1.806919 | -0.017770 |
| H | 2.611593 | 2.450216 | -0.374012 | N | 0.537232 | -1.604684 | 0.102258 |
| H | 2.981907 | -2.491793 | 0.335197 | N | 0.319618 | 1.162596 | -0.316187 |
| H | 5.036410 | -1.075598 | 0.235830 | O | -1.784327 | -3.076709 | 0.212221 |
| C | 1.539267 | 0.583260 | -0.163959 | H | -2.102076 | 1.502642 | -1.422926 |
| C | 1.642024 | -0.822280 | 0.039668 | O | -1.319712 | 3.083056 | 0.521617 |
| C | -0.743923 | 0.391314 | -0.239791 | N | -2.317559 | 2.185423 | 0.544159 |
| C | -2.083529 | 1.072137 | -0.399879 | H | -0.520425 | 2.685405 | 0.085659 |
| C | -3.251923 | 0.143673 | -0.274097 | | | | |

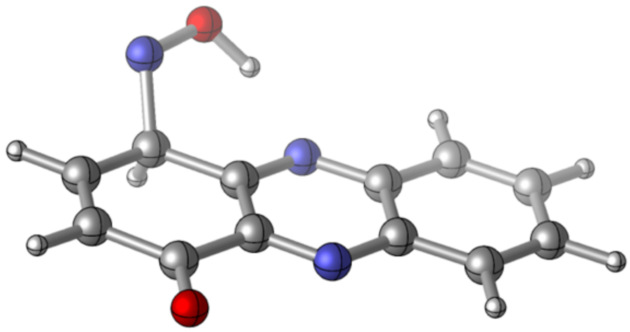

Zero-point correction= 0.184243 (Hartree/Particle)
Thermal correction to Energy= 0.196547
Thermal correction to Enthalpy= 0.197491
Thermal correction to Gibbs Free Energy= 0.144474
Sum of electronic and zero-point Energies= -776.312549
Sum of electronic and thermal Energies= -776.300244
Sum of electronic and thermal Enthalpies= -776.299300
Sum of electronic and thermal Free Energies= -776.352317

**$^{4}$TS$_{3'\text{-}4}$**

E(scf) = -906.335507352 a.u.
$\nu_{min}$ = -2059.3 $cm^{-1}$

| | | | |
|---|---|---|---|
| C | -2.136858 | 2.465421 | -0.034217 |
| C | -3.384497 | 1.892953 | -0.053360 |
| C | -3.077008 | 4.702402 | -0.048475 |
| C | -1.980691 | 3.878135 | -0.030240 |
| H | -1.249512 | 1.831152 | -0.020229 |
| H | -3.522165 | 0.811223 | -0.054012 |
| H | -2.985002 | 5.788917 | -0.047535 |
| H | -0.977051 | 4.304506 | -0.012942 |
| C | -4.532296 | 2.724792 | -0.070902 |
| C | -4.381818 | 4.140672 | -0.072275 |
| C | -6.813386 | 2.980804 | -0.109685 |
| C | -8.178322 | 2.378789 | -0.072191 |
| C | -9.294403 | 3.321790 | -0.287332 |
| C | -9.156728 | 4.659339 | -0.291323 |
| C | -7.848456 | 5.314847 | -0.168542 |
| C | -6.649373 | 4.400990 | -0.119546 |
| H | -10.276664 | 2.860435 | -0.408379 |
| H | -10.019558 | 5.316626 | -0.406644 |
| N | -5.460561 | 4.960663 | -0.095899 |
| N | -5.771641 | 2.172634 | -0.072721 |
| O | -7.717080 | 6.525409 | -0.128514 |
| H | -8.240519 | 2.216580 | 1.263613 |
| O | -7.462193 | 0.244586 | -0.599541 |
| N | -8.470042 | 1.132213 | -0.609359 |
| N | -8.092480 | 2.229669 | 2.630106 |

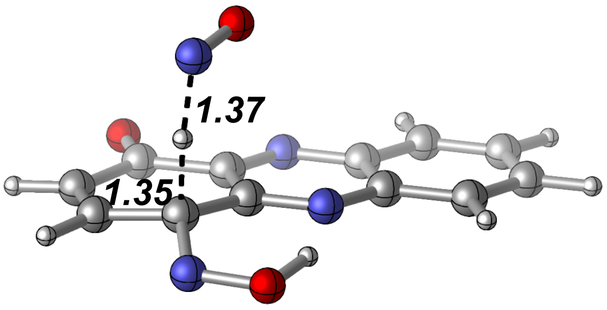


Zero-point correction= 0.185116 (Hartree/Particle)
Thermal correction to Energy= 0.200161
Thermal correction to Enthalpy= 0.201105
Thermal correction to Gibbs Free Energy= 0.142050
Sum of electronic and zero-point Energies= -906.150392
Sum of electronic and thermal Energies= -906.135347
Sum of electronic and thermal Enthalpies= -906.134402
Sum of electronic and thermal Free Energies= -906.193458

[4]**4**

E(scf) = -775.936140550 a.u.

| | | | | | | | |
|---|---|---|---|---|---|---|---|
| C | 3.923677 | 0.785467 | 0.000112 | C | -3.263602 | 0.150702 | -0.000041 |
| C | 2.694082 | 1.396317 | 0.000148 | C | -3.170899 | -1.192517 | -0.000055 |
| C | 2.914090 | -1.421560 | -0.000139 | C | -1.877370 | -1.888988 | 0.000041 |
| C | 4.035631 | -0.631020 | -0.000036 | C | -0.646577 | -1.013909 | -0.000119 |
| H | 4.830302 | 1.391923 | 0.000197 | H | -4.234536 | 0.649182 | -0.000027 |
| H | 2.589932 | 2.481755 | 0.000256 | H | -4.061046 | -1.822476 | -0.000039 |
| H | 2.972837 | -2.510323 | -0.000237 | N | 0.523880 | -1.608801 | -0.000129 |
| H | 5.025718 | -1.088322 | -0.000057 | N | 0.299976 | 1.190380 | 0.000041 |
| C | 1.521570 | 0.599941 | 0.000024 | O | -1.781765 | -3.103272 | 0.000326 |
| C | 1.627646 | -0.820406 | -0.000108 | N | -2.368108 | 2.304546 | 0.000015 |
| C | -0.766911 | 0.411472 | -0.000150 | O | -1.363258 | 3.177553 | 0.000026 |
| C | -2.100542 | 1.028574 | -0.000103 | H | -0.498005 | 2.676704 | 0.000169 |

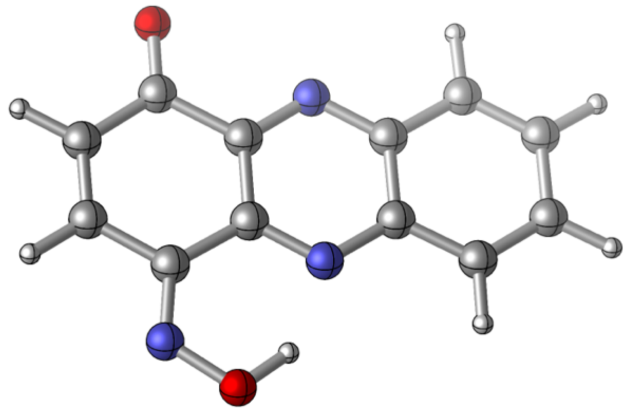

Zero-point correction= 0.174218 (Hartree/Particle)
Thermal correction to Energy= 0.185949
Thermal correction to Enthalpy= 0.186894
Thermal correction to Gibbs Free Energy= 0.135976
Sum of electronic and zero-point Energies= -775.761922
Sum of electronic and thermal Energies= -775.750191
Sum of electronic and thermal Enthalpies= -775.749247
Sum of electronic and thermal Free Energies= -775.800165

**$^{2}Int_{I}$**
E(scf) = -776.474903506 a.u.

| | | | | | | | |
|---|---|---|---|---|---|---|---|
| C | -4.459115 | -0.487149 | 0.243972 | C | 2.606307 | -1.425494 | -0.417596 |
| C | -3.357613 | -1.312553 | 0.141412 | C | 2.874070 | 0.039327 | -0.481947 |
| C | -3.047279 | 1.479329 | 0.112156 | C | 1.669829 | 0.911837 | -0.340160 |
| C | -4.299679 | 0.918494 | 0.229146 | C | 0.369255 | 0.393229 | -0.210416 |
| H | -5.457436 | -0.917229 | 0.337191 | H | 1.182569 | -2.991651 | -0.251129 |
| H | -3.454467 | -2.399524 | 0.150838 | H | 3.469702 | -2.087924 | -0.479972 |
| H | -2.902584 | 2.560826 | 0.098514 | N | -0.674720 | 1.232089 | -0.109762 |
| H | -5.178027 | 1.560916 | 0.311251 | N | -0.983544 | -1.606075 | -0.078011 |
| C | -2.063973 | -0.761766 | 0.020712 | N | 3.888984 | 0.557945 | 0.546408 |
| C | -1.897153 | 0.653409 | 0.004529 | O | 4.507529 | -0.288583 | 1.107787 |
| C | 0.192994 | -1.050269 | -0.191842 | H | 3.398448 | 0.299922 | -1.425726 |
| C | 1.368581 | -1.917047 | -0.287764 | O | 1.842702 | 2.238588 | -0.347880 |

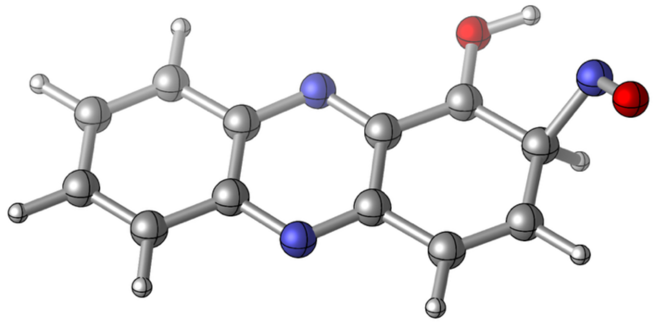

Zero-point correction= 0.182600 (Hartree/Particle)
Thermal correction to Energy= 0.195451
Thermal correction to Enthalpy= 0.196395
Thermal correction to Gibbs Free Energy= 0.141748
Sum of electronic and zero-point Energies= -776.292304
Sum of electronic and thermal Energies= -776.279453
Sum of electronic and thermal Enthalpies= -776.278509
Sum of electronic and thermal Free Energies= -776.333155

**$^{2}TS_{I}$**
E(scf) = -906.321865461 a.u.
$\nu_{min}$ = -1879.8cm$^{-1}$

| | | | | | | | |
|---|---|---|---|---|---|---|---|
| C | -0.634628 | -0.631292 | 0.199388 | C | -7.011569 | 2.411096 | -0.932832 |
| C | -1.440493 | 0.484208 | 0.186085 | C | -7.670579 | 1.117500 | -1.279044 |
| C | -2.466687 | -2.031191 | -0.551719 | C | -6.764190 | -0.080113 | -1.303067 |
| C | -1.151748 | -1.897711 | -0.171923 | C | -5.361586 | 0.045005 | -0.939251 |
| H | 0.411308 | -0.544571 | 0.497291 | H | -5.265232 | 3.458781 | -0.333190 |
| H | -1.063902 | 1.468647 | 0.467612 | H | -7.640670 | 3.301704 | -0.942152 |
| H | -2.887023 | -2.995171 | -0.841912 | N | -4.617673 | -1.045768 | -0.949527 |
| H | -0.497838 | -2.770867 | -0.155111 | N | -3.584436 | 1.491758 | -0.206113 |
| C | -2.798571 | 0.373678 | -0.198721 | N | -8.832126 | 0.746150 | -0.332661 |
| C | -3.318673 | -0.897322 | -0.572639 | O | -9.128891 | 1.606633 | 0.433774 |
| C | -4.836982 | 1.333834 | -0.562999 | H | -8.196248 | 1.178918 | -2.249219 |
| C | -5.719829 | 2.501459 | -0.593458 | O | -7.220281 | -1.192451 | -1.769884 |

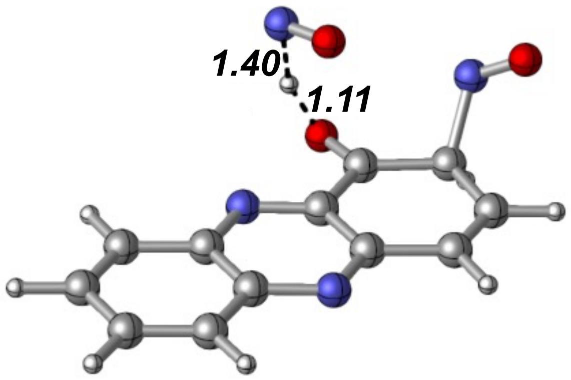


Zero-point correction= 0.184762 (Hartree/Particle)
Thermal correction to Energy= 0.199745
Thermal correction to Enthalpy= 0.200689
Thermal correction to Gibbs Free Energy= 0.141467
Sum of electronic and zero-point Energies= -906.137103
Sum of electronic and thermal Energies= -906.122120
Sum of electronic and thermal Enthalpies= -906.121176
Sum of electronic and thermal Free Energies= -906.180398

**$^{2}$INT$_{II}$**
E(scf) = -775.893271057 a.u.

| | | | | | | | |
|---|---|---|---|---|---|---|---|
| C | -4.412941 | -0.515433 | 0.270029 | C | 2.652383 | -1.350757 | -0.463265 |
| C | -3.316845 | -1.334215 | 0.150331 | C | 2.922986 | 0.115035 | -0.473067 |
| C | -3.032577 | 1.473418 | 0.128503 | C | 1.721705 | 1.051684 | -0.380807 |
| C | -4.271655 | 0.898195 | 0.259291 | C | 0.367175 | 0.419636 | -0.236033 |
| H | -5.407548 | -0.951351 | 0.375046 | H | 1.242124 | -2.933655 | -0.349264 |
| H | -3.405505 | -2.421182 | 0.156429 | H | 3.515490 | -2.012359 | -0.543705 |
| H | -2.895437 | 2.555340 | 0.116984 | N | -0.660957 | 1.228710 | -0.125697 |
| H | -5.158585 | 1.525663 | 0.356028 | N | -0.944259 | -1.585895 | -0.103229 |
| C | -2.023010 | -0.767350 | 0.014263 | O | 1.874485 | 2.251310 | -0.411365 |
| C | -1.881692 | 0.650634 | 0.003340 | N | 3.900744 | 0.584739 | 0.612506 |
| C | 0.233313 | -1.006392 | -0.225906 | O | 4.413135 | -0.305264 | 1.213352 |
| C | 1.419687 | -1.857188 | -0.350896 | H | 3.468621 | 0.412698 | -1.387177 |

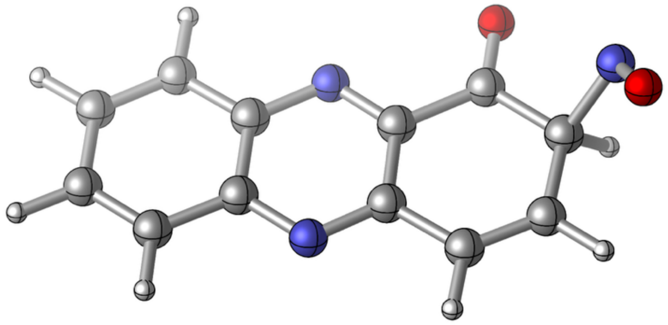

Zero-point correction= 0.171862 (Hartree/Particle)
Thermal correction to Energy= 0.184270
Thermal correction to Enthalpy= 0.185214
Thermal correction to Gibbs Free Energy= 0.131822
Sum of electronic and zero-point Energies= -775.721409
Sum of electronic and thermal Energies= -775.709001
Sum of electronic and thermal Enthalpies= -775.708057
Sum of electronic and thermal Free Energies= -775.761449

**$^2TS_{II}$**
E(scf) = -852.286628227 a.u.
$\nu_{min}$ = -103.7 $cm^{-1}$

| | | | | | | | |
|---|---|---|---|---|---|---|---|
| C | -1.672812 | -1.882186 | -0.110363 | C | 5.356557 | -2.562200 | 0.926547 |
| C | -0.585747 | -2.677395 | 0.151576 | C | 5.583055 | -1.145189 | 0.679240 |
| C | -0.288060 | 0.107238 | -0.177919 | C | 4.454525 | -0.215054 | 0.540702 |
| C | -1.524804 | -0.477556 | -0.276751 | C | 3.103152 | -0.882895 | 0.429023 |
| H | -2.665462 | -2.327725 | -0.193167 | H | 3.968181 | -4.182845 | 1.061410 |
| H | -0.679159 | -3.756421 | 0.282046 | H | 6.242296 | -3.159977 | 1.147391 |
| H | -0.145888 | 1.181858 | -0.300825 | N | 2.072750 | -0.101173 | 0.187726 |
| H | -2.404700 | 0.133105 | -0.484063 | N | 1.775831 | -2.894256 | 0.515849 |
| C | 0.709339 | -2.099215 | 0.258382 | O | 4.557722 | 1.002316 | 0.477099 |
| C | 0.856810 | -0.691953 | 0.091918 | N | 6.827466 | -0.562774 | 0.888369 |
| C | 2.960201 | -2.305599 | 0.596672 | O | 7.785619 | -1.342951 | 0.971296 |
| C | 4.128998 | -3.119943 | 0.879384 | H | 6.065856 | -1.382650 | -0.888195 |

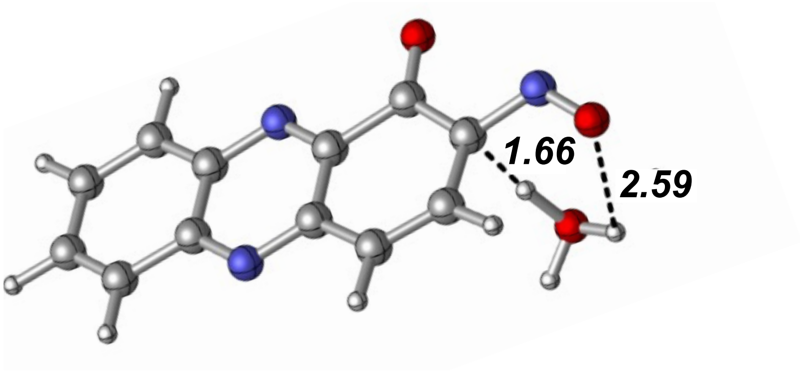


Zero-point correction= 0.195228 (Hartree/Particle)
Thermal correction to Energy= 0.209597
Thermal correction to Enthalpy= 0.210541
Thermal correction to Gibbs Free Energy= 0.153745
Sum of electronic and zero-point Energies= -852.091400
Sum of electronic and thermal Energies= -852.077031
Sum of electronic and thermal Enthalpies= -852.076087
Sum of electronic and thermal Free Energies= -852.132884

**$^{4}INT_{I}$**

E(scf) = -776.480081245 a.u.

| | | | | | | | |
|---|---|---|---|---|---|---|---|
| C | -4.065649 | -0.958615 | -0.071913 | C | 3.090012 | -0.216745 | -0.687578 |
| C | -2.832372 | -1.515539 | -0.327921 | C | 2.945010 | 1.110024 | -0.404773 |
| C | -3.052512 | 1.200071 | 0.373916 | C | 1.694779 | 1.677914 | -0.052874 |
| C | -4.174078 | 0.408823 | 0.283093 | C | 0.499763 | 0.880822 | -0.060915 |
| H | -4.966356 | -1.570387 | -0.140512 | H | 4.067477 | -0.615779 | -0.955753 |
| H | -2.721823 | -2.566475 | -0.599582 | H | 3.813035 | 1.772378 | -0.463696 |
| H | -3.114305 | 2.255725 | 0.642525 | N | -0.673865 | 1.451590 | 0.188714 |
| H | -5.158342 | 0.834826 | 0.484261 | N | -0.444295 | -1.294555 | -0.480564 |
| C | -1.664653 | -0.722834 | -0.237120 | N | 2.101185 | -2.145635 | 0.646104 |
| C | -1.769405 | 0.651583 | 0.111641 | O | 3.139106 | -2.061627 | 1.219367 |
| C | 0.604535 | -0.523510 | -0.383381 | H | 1.900946 | -1.885149 | -1.389900 |
| C | 1.956768 | -1.165398 | -0.556136 | O | 1.547976 | 2.976522 | 0.256418 |

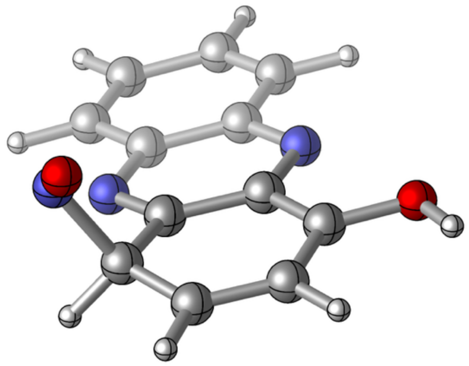

Zero-point correction= 0.182368 (Hartree/Particle)
Thermal correction to Energy= 0.195201
Thermal correction to Enthalpy= 0.196145
Thermal correction to Gibbs Free Energy= 0.141562
Sum of electronic and zero-point Energies= -776.297713
Sum of electronic and thermal Energies= -776.284880
Sum of electronic and thermal Enthalpies= -776.283936
Sum of electronic and thermal Free Energies= -776.338519

**$^4TS_I$**

E(scf) = -906.323952492 a.u.

$\nu_{min}$ = -1952.0 $cm^{-1}$

| | | | | | | | |
|---|---|---|---|---|---|---|---|
| C | -0.352306 | 0.032038 | 0.023783 | C | 6.753465 | -0.392536 | -1.224921 |
| C | 0.768730 | -0.741495 | -0.163267 | C | 6.814793 | 0.952114 | -1.269774 |
| C | 0.947126 | 2.070550 | -0.182800 | C | 5.664344 | 1.788448 | -0.998229 |
| C | -0.261540 | 1.447971 | 0.015515 | C | 4.364288 | 1.150949 | -0.755226 |
| H | -1.322112 | -0.442189 | 0.181214 | H | 7.639759 | -0.989255 | -1.442092 |
| H | 0.723955 | -1.831392 | -0.157947 | H | 7.738606 | 1.467770 | -1.537774 |
| H | 1.040236 | 3.157314 | -0.196335 | N | 3.307622 | 1.921840 | -0.589912 |
| H | -1.163005 | 2.043532 | 0.167233 | N | 3.136407 | -0.901077 | -0.534624 |
| C | 2.028049 | -0.123824 | -0.364503 | N | 5.655035 | -1.842087 | 0.543499 |
| C | 2.119909 | 1.295374 | -0.380242 | O | 6.740769 | -1.770242 | 1.024665 |
| C | 4.275593 | -0.280849 | -0.711684 | H | 5.306025 | -1.978685 | -1.484272 |
| C | 5.520421 | -1.124041 | -0.820711 | O | 5.733796 | 3.072852 | -1.137769 |

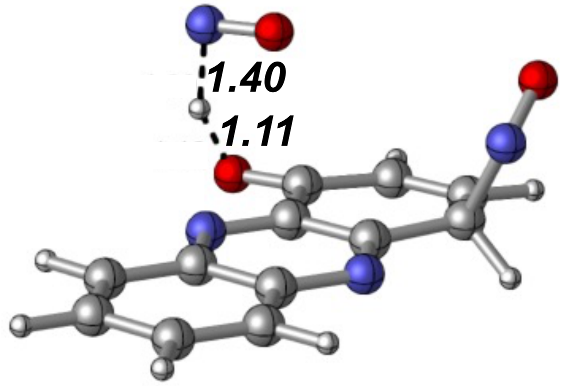


Zero-point correction= 0.184468 (Hartree/Particle)
Thermal correction to Energy= 0.199596
Thermal correction to Enthalpy= 0.200540
Thermal correction to Gibbs Free Energy= 0.140464
Sum of electronic and zero-point Energies= -906.139484
Sum of electronic and thermal Energies= -906.124357
Sum of electronic and thermal Enthalpies= -906.123413
Sum of electronic and thermal Free Energies= -906.183489

**$^{4}$INT$_{II}$**
E(scf) = -775.894727217 a.u.

| | | | | | | | |
|---|---|---|---|---|---|---|---|
| C | -4.019066 | -1.027035 | -0.052834 | C | 3.126489 | -0.067185 | -0.544275 |
| C | -2.774714 | -1.562291 | -0.273925 | C | 2.978030 | 1.240996 | -0.301365 |
| C | -3.083606 | 1.180998 | 0.318432 | C | 1.668056 | 1.853877 | -0.006039 |
| C | -4.175455 | 0.353995 | 0.246140 | C | 0.471568 | 0.938806 | -0.050084 |
| H | -4.902721 | -1.664600 | -0.105976 | H | 4.119772 | -0.476774 | -0.734486 |
| H | -2.632967 | -2.618858 | -0.503909 | H | 3.832114 | 1.920444 | -0.303517 |
| H | -3.176871 | 2.243922 | 0.544095 | N | -0.709249 | 1.479216 | 0.158866 |
| H | -5.175768 | 0.753766 | 0.417513 | N | -0.397563 | -1.265410 | -0.419892 |
| C | -1.626751 | -0.730289 | -0.202145 | O | 1.551798 | 3.035878 | 0.255551 |
| C | -1.782423 | 0.654316 | 0.094366 | N | 2.158758 | -2.159052 | 0.498641 |
| C | 0.628418 | -0.449037 | -0.337205 | O | 3.215911 | -2.161137 | 1.045950 |
| C | 1.997629 | -1.040072 | -0.542390 | H | 1.982080 | -1.621583 | -1.481075 |

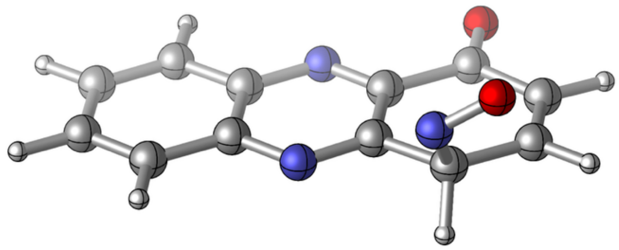

Zero-point correction= 0.172048 (Hartree/Particle)
Thermal correction to Energy= 0.184499
Thermal correction to Enthalpy= 0.185443
Thermal correction to Gibbs Free Energy= 0.131980
Sum of electronic and zero-point Energies= -775.722679
Sum of electronic and thermal Energies= -775.710228
Sum of electronic and thermal Enthalpies= -775.709284
Sum of electronic and thermal Free Energies= -775.762747

**$^{4}TS_{II}$**
E(scf) = -852.284484010 a.u.
$\nu_{min}$ = -120.4 $cm^{-1}$

| | | | | | | | |
|---|---|---|---|---|---|---|---|
| C | -1.273977 | -0.515190 | -0.046988 | C | 5.551576 | 1.677118 | 0.646873 |
| C | 0.043008 | -0.834368 | 0.165093 | C | 5.184402 | 2.965651 | 0.445814 |
| C | -0.739424 | 1.838415 | -0.277961 | C | 3.804500 | 3.338019 | 0.155893 |
| C | -1.671098 | 0.832745 | -0.271055 | C | 2.803298 | 2.210397 | 0.147230 |
| H | -2.031112 | -1.301088 | -0.045493 | H | 6.585455 | 1.422291 | 0.888646 |
| H | 0.366205 | -1.861922 | 0.337596 | H | 5.909362 | 3.778092 | 0.517008 |
| H | -1.015242 | 2.880415 | -0.446609 | N | 1.544989 | 2.537716 | -0.065862 |
| H | -2.725000 | 1.060192 | -0.437470 | N | 2.329359 | -0.141291 | 0.367341 |
| C | 1.032975 | 0.188108 | 0.164358 | O | 3.454374 | 4.493970 | -0.050520 |
| C | 0.634075 | 1.537171 | -0.058839 | N | 4.986228 | -0.705965 | 0.997317 |
| C | 3.209653 | 0.847624 | 0.370116 | O | 6.199807 | -0.924809 | 1.117166 |
| C | 4.635034 | 0.563230 | 0.556117 | H | 5.158761 | 0.108802 | -0.923188 |

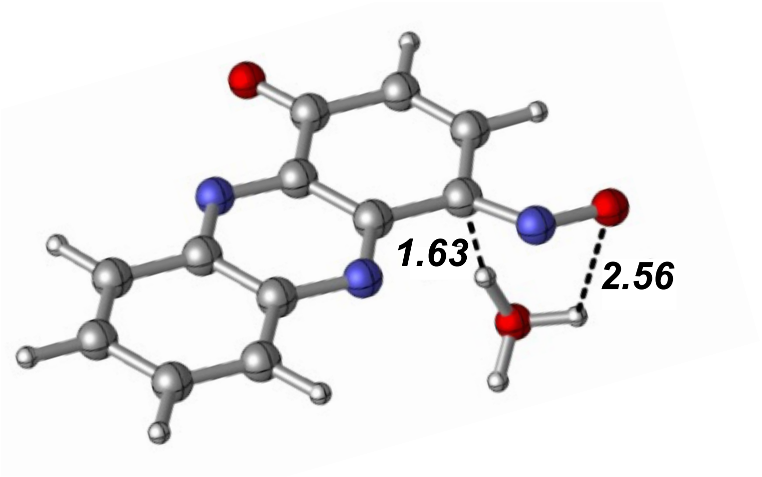


Zero-point correction= 0.194771 (Hartree/Particle)
Thermal correction to Energy= 0.209242
Thermal correction to Enthalpy= 0.210186
Thermal correction to Gibbs Free Energy= 0.153114
Sum of electronic and zero-point Energies= -852.089713
Sum of electronic and thermal Energies= -852.075242
Sum of electronic and thermal Enthalpies= -852.074298
Sum of electronic and thermal Free Energies= -852.131370

**$^2TS_{2\text{-}2'}$**
E(scf) = -982.785172750 a.u.
$\nu_{min}$ = -171.3 $cm^{-1}$

| | | | | | | | |
|---|---|---|---|---|---|---|---|
| C | -1.540838 | 1.707710 | 0.286143 | C | 5.134335 | -0.502861 | -1.025839 |
| C | -0.644781 | 0.690595 | 0.065234 | C | 5.698226 | 0.882843 | -0.984629 |
| C | 0.198354 | 3.385339 | 0.079015 | C | 4.696935 | 1.992384 | -0.771086 |
| C | -1.118345 | 3.064255 | 0.293005 | C | 3.274641 | 1.679092 | -0.551138 |
| H | -2.592887 | 1.476245 | 0.458740 | H | 3.445765 | -1.773419 | -0.840186 |
| H | -0.951077 | -0.355921 | 0.055307 | H | 5.846268 | -1.309710 | -1.204458 |
| H | 0.549923 | 4.417647 | 0.079738 | N | 2.446856 | 2.678820 | -0.354990 |
| H | -1.851286 | 3.852133 | 0.470733 | N | 1.601298 | -0.022441 | -0.380049 |
| C | 0.723241 | 0.991536 | -0.158478 | O | 4.994922 | 3.203335 | -1.084966 |
| C | 1.146937 | 2.353523 | -0.150206 | H | 6.330822 | 1.092224 | -1.857389 |
| C | 2.861226 | 0.308361 | -0.572807 | N | 6.525649 | 1.119827 | 0.232737 |
| C | 3.836894 | -0.755500 | -0.824724 | O | 7.798359 | 1.095850 | 0.285892 |

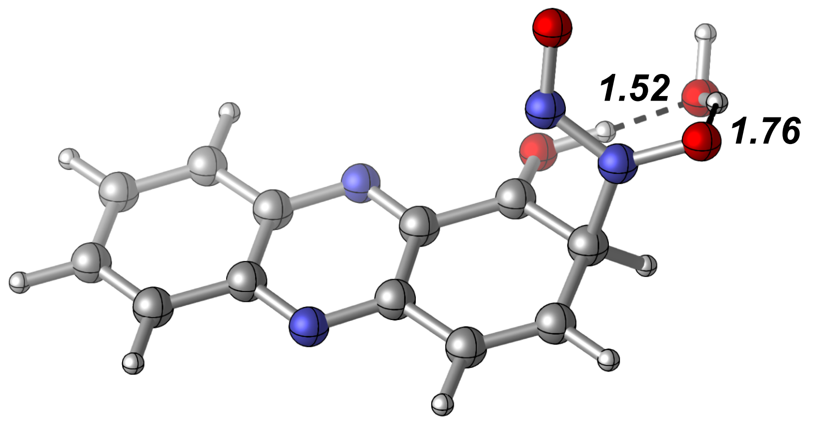


Zero-point correction= 0.218554 (Hartree/Particle)
Thermal correction to Energy= 0.234248
Thermal correction to Enthalpy= 0.235193
Thermal correction to Gibbs Free Energy= 0.174908
Sum of electronic and zero-point Energies= -982.566619
Sum of electronic and thermal Energies= -982.550924
Sum of electronic and thermal Enthalpies= -982.549980
Sum of electronic and thermal Free Energies= -982.610264

**$^2$2'**

E(scf) = -906.401218810 a.u.

| | | | | | | | |
|---|---|---|---|---|---|---|---|
| C | 5.070767 | 0.164540 | 0.271800 | C | -1.857921 | 1.789542 | -0.424697 |
| C | 4.068146 | 1.101032 | 0.213966 | C | -2.226749 | 0.372506 | -0.753976 |
| C | 3.484058 | -1.656402 | 0.055368 | C | -1.200366 | -0.675973 | -0.314559 |
| C | 4.778736 | -1.223906 | 0.192691 | C | 0.219748 | -0.223361 | -0.211807 |
| H | 6.107769 | 0.485650 | 0.380405 | H | -0.325305 | 3.194819 | -0.014890 |
| H | 4.273583 | 2.170381 | 0.273420 | H | -2.660506 | 2.526380 | -0.471501 |
| H | 3.231168 | -2.715414 | -0.006953 | N | 1.149804 | -1.147269 | -0.138181 |
| H | 5.594402 | -1.946248 | 0.242742 | N | 1.734559 | 1.619368 | 0.018278 |
| C | 2.719167 | 0.684073 | 0.071211 | O | -1.547536 | -1.812607 | -0.068154 |
| C | 2.426918 | -0.709835 | -0.008031 | H | -2.288874 | 0.289304 | -1.854833 |
| C | 0.501353 | 1.178383 | -0.130911 | N | -3.547660 | 0.009765 | -0.261619 |
| C | -0.594556 | 2.151016 | -0.179057 | O | -4.121629 | -1.095331 | -0.829961 |

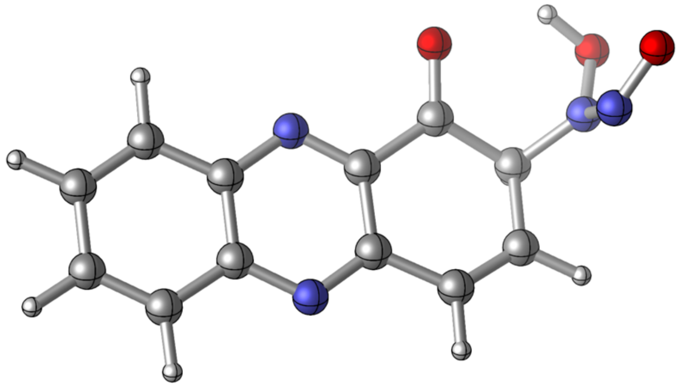

Zero-point correction= 0.193871 (Hartree/Particle)
Thermal correction to Energy= 0.208162
Thermal correction to Enthalpy= 0.209106
Thermal correction to Gibbs Free Energy= 0.151701
Sum of electronic and zero-point Energies= -906.207348
Sum of electronic and thermal Energies= -906.193057
Sum of electronic and thermal Enthalpies= -906.192113
Sum of electronic and thermal Free Energies= -906.249518

**$^{2}TS_{2'-3*}$**
E(scf) = -982.788519057 a.u.
$\nu_{min}$ = -395.3 cm$^{-1}$

| | | | | | | | |
|---|---|---|---|---|---|---|---|
| C | -0.622591 | 0.262453 | 0.070185 | C | -7.518474 | 1.987799 | 0.658513 |
| C | -1.595516 | 1.221904 | 0.167548 | C | -7.891006 | 0.598408 | 0.504509 |
| C | -2.263078 | -1.520132 | 0.197099 | C | -6.920094 | -0.450366 | 0.648495 |
| C | -0.957454 | -1.122628 | 0.084894 | C | -5.503478 | -0.011658 | 0.495327 |
| H | 0.424470 | 0.556232 | -0.019914 | H | -5.932950 | 3.430448 | 0.642252 |
| H | -1.358578 | 2.286695 | 0.157858 | H | -8.322580 | 2.721097 | 0.748862 |
| H | -2.547623 | -2.573205 | 0.211743 | N | -4.580818 | -0.952890 | 0.415956 |
| H | -0.162789 | -1.865888 | 0.006350 | N | -3.912334 | 1.797320 | 0.380180 |
| C | -2.964537 | 0.841835 | 0.284815 | O | -7.232587 | -1.631987 | 0.890664 |
| C | -3.298583 | -0.546724 | 0.300750 | H | -7.995532 | 0.669633 | -1.139496 |
| C | -5.176207 | 1.392207 | 0.481117 | N | -9.231324 | 0.181397 | 0.816956 |
| C | -6.221924 | 2.381025 | 0.601312 | O | -9.429283 | -0.581563 | 1.935615 |

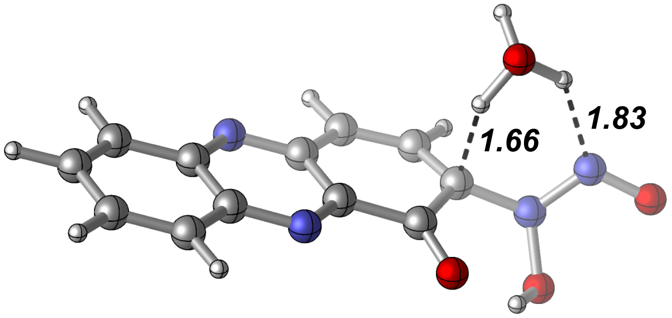


Zero-point correction= 0.216389 (Hartree/Particle)
Thermal correction to Energy= 0.232266
Thermal correction to Enthalpy= 0.233210
Thermal correction to Gibbs Free Energy= 0.172749
Sum of electronic and zero-point Energies= -982.572130
Sum of electronic and thermal Energies= -982.556253
Sum of electronic and thermal Enthalpies= -982.555309
Sum of electronic and thermal Free Energies= -982.615770

**²3***

E(scf) = -906.366237579 a.u.

| | | | | | | | |
|---|---|---|---|---|---|---|---|
| C | 5.182693 | 0.279922 | 0.071369 | C | -1.821818 | 1.606979 | -0.198800 |
| C | 4.154509 | 1.182779 | -0.003691 | C | -2.106907 | 0.208728 | -0.127143 |
| C | 3.635726 | -1.587246 | 0.127523 | C | -1.094295 | -0.817536 | -0.033165 |
| C | 4.923210 | -1.119136 | 0.138033 | C | 0.303470 | -0.267614 | -0.034357 |
| H | 6.215867 | 0.631171 | 0.080489 | H | -0.323745 | 3.135069 | -0.271442 |
| H | 4.333953 | 2.257632 | -0.055302 | H | -2.649034 | 2.312596 | -0.310663 |
| H | 3.407952 | -2.653049 | 0.177265 | N | 1.281023 | -1.150561 | 0.038888 |
| H | 5.759933 | -1.816646 | 0.197527 | N | 1.799731 | 1.626782 | -0.088562 |
| C | 2.803841 | 0.727782 | -0.015790 | O | -1.316853 | -2.023722 | 0.068746 |
| C | 2.545206 | -0.674252 | 0.049940 | N | -3.435335 | -0.192058 | -0.072486 |
| C | 0.557058 | 1.149640 | -0.095816 | O | -3.885229 | -1.222097 | -0.789218 |
| C | -0.545492 | 2.072651 | -0.182684 | N | -4.360454 | 0.404949 | 0.606986 |

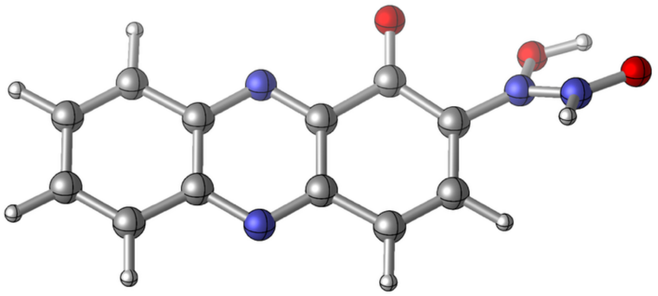

Zero-point correction= 0.194521 (Hartree/Particle)
Thermal correction to Energy= 0.208643
Thermal correction to Enthalpy= 0.209587
Thermal correction to Gibbs Free Energy= 0.152643
Sum of electronic and zero-point Energies= -906.171717
Sum of electronic and thermal Energies= -906.157595
Sum of electronic and thermal Enthalpies= -906.156651
Sum of electronic and thermal Free Energies= -906.213594

**$^{2}$TS$_{3*-4'}$**
E(scf) = -906.353450848 a.u.
$\nu_{min}$ = -544.5 cm$^{-1}$

| | | | | | | | |
|---|---|---|---|---|---|---|---|
| C | -1.711142 | 1.669053 | 0.004692 | C | 5.266372 | 3.084425 | -0.517464 |
| C | -0.692779 | 2.582689 | -0.072501 | C | 5.581482 | 1.678761 | -0.524451 |
| C | -0.154490 | -0.190329 | -0.089784 | C | 4.546743 | 0.674281 | -0.413360 |
| C | -1.441420 | 0.270303 | -0.004252 | C | 3.151778 | 1.168741 | -0.330656 |
| H | -2.744847 | 2.011235 | 0.073802 | H | 3.744338 | 4.580302 | -0.418134 |
| H | -0.879400 | 3.657223 | -0.067389 | H | 6.084746 | 3.798948 | -0.615591 |
| H | 0.079689 | -1.255517 | -0.098529 | N | 2.190531 | 0.267452 | -0.255563 |
| H | -2.271909 | -0.434171 | 0.057737 | N | 1.651823 | 3.047236 | -0.240834 |
| C | 0.656392 | 2.137090 | -0.163231 | O | 4.809369 | -0.547953 | -0.376703 |
| C | 0.926481 | 0.733331 | -0.171248 | N | 6.879354 | 1.322335 | -0.628966 |
| C | 2.890272 | 2.578899 | -0.323738 | O | 7.178584 | 0.041304 | -0.697651 |
| C | 3.989766 | 3.518706 | -0.416233 | N | 8.052145 | 2.001219 | 0.563022 |

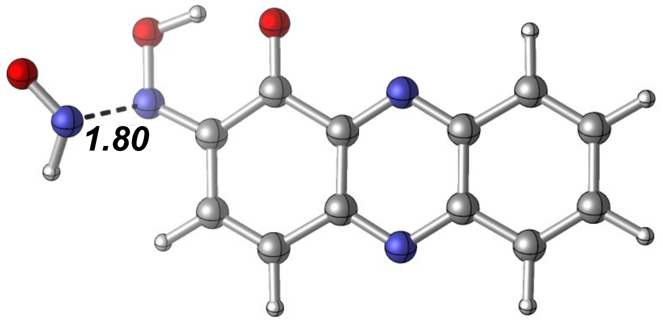


Zero-point correction= 0.190274 (Hartree/Particle)
Thermal correction to Energy= 0.204344
Thermal correction to Enthalpy= 0.205288
Thermal correction to Gibbs Free Energy= 0.148539
Sum of electronic and zero-point Energies= -906.163177
Sum of electronic and thermal Energies= -906.149107
Sum of electronic and thermal Enthalpies= -906.148163
Sum of electronic and thermal Free Energies= -906.204911

**$^4$TS$_{2\text{-}2'}$**
E(scf) = -982.786817815 a.u.
$\nu_{min}$ = -761.9 $cm^{-1}$

| | | | | | | | |
|---|---|---|---|---|---|---|---|
| C | -0.292538 | -0.201623 | 0.114567 | C | -7.081494 | 0.713843 | 1.890562 |
| C | -1.470143 | -0.747007 | 0.569094 | C | -7.042872 | 1.990791 | 1.467811 |
| C | -1.414708 | 1.851736 | -0.521514 | C | -6.135375 | 2.302699 | 0.366809 |
| C | -0.264591 | 1.104275 | -0.436534 | C | -4.858650 | 1.516648 | 0.333993 |
| H | 0.631830 | -0.777816 | 0.173732 | H | -7.790922 | 0.378841 | 2.647721 |
| H | -1.514503 | -1.752049 | 0.990101 | H | -7.715574 | 2.768949 | 1.828915 |
| H | -1.418094 | 2.860002 | -0.937187 | N | -3.772832 | 2.075133 | -0.131432 |
| H | 0.680345 | 1.515555 | -0.793925 | N | -3.841234 | -0.571579 | 0.913250 |
| C | -2.670564 | -0.000446 | 0.490736 | O | -6.302587 | 3.283915 | -0.399093 |
| C | -2.640128 | 1.315624 | -0.053722 | O | -6.953250 | 0.603695 | -0.962198 |
| C | -4.899376 | 0.185697 | 0.837448 | N | -6.954082 | -0.460520 | -0.186310 |
| C | -6.280251 | -0.318925 | 1.139413 | N | -7.874188 | -1.412044 | -0.246904 |

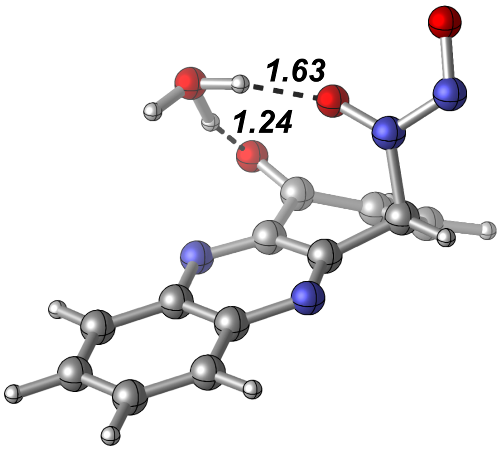


Zero-point correction= 0.215453 (Hartree/Particle)
Thermal correction to Energy= 0.230840
Thermal correction to Enthalpy= 0.231785
Thermal correction to Gibbs Free Energy= 0.172806
Sum of electronic and zero-point Energies= -982.571364
Sum of electronic and thermal Energies= -982.555977
Sum of electronic and thermal Enthalpies= -982.555033
Sum of electronic and thermal Free Energies= -982.614012

**[4]2'**

E(scf) = -906.404990314 a.u.

| | | | | | | | |
|---|---|---|---|---|---|---|---|
| C | -3.889914 | -1.752253 | -0.046704 | C | 2.696372 | 1.189666 | -0.462607 |
| C | -2.553628 | -1.901314 | -0.321194 | C | 2.194517 | 2.379308 | -0.116392 |
| C | -3.613595 | 0.620012 | 0.386187 | C | 0.749346 | 2.617381 | 0.065941 |
| C | -4.424312 | -0.483284 | 0.308663 | C | -0.149981 | 1.416616 | -0.069568 |
| H | -4.555592 | -2.614806 | -0.100620 | H | 3.772675 | 1.052631 | -0.583188 |
| H | -2.123787 | -2.864863 | -0.596800 | H | 2.846535 | 3.238035 | 0.049658 |
| H | -3.998438 | 1.603410 | 0.658367 | N | -1.427618 | 1.587070 | 0.195360 |
| H | -5.490391 | -0.393622 | 0.521318 | N | -0.372164 | -0.923519 | -0.532778 |
| C | -1.691723 | -0.775763 | -0.250750 | O | 0.304198 | 3.715886 | 0.334461 |
| C | -2.224897 | 0.495945 | 0.108412 | H | 1.934920 | -0.198297 | -1.864004 |
| C | 0.377349 | 0.148242 | -0.443431 | O | 2.391265 | -2.353444 | -0.868527 |
| C | 1.850488 | -0.008333 | -0.781584 | N | 2.416652 | -1.195946 | -0.162262 |

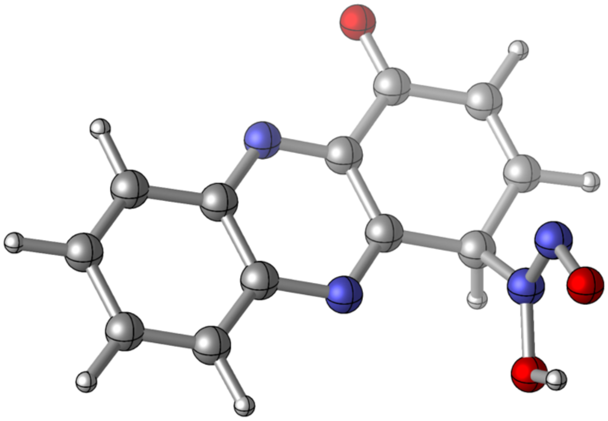

Zero-point correction= 0.194034 (Hartree/Particle)
Thermal correction to Energy= 0.208402
Thermal correction to Enthalpy= 0.209346
Thermal correction to Gibbs Free Energy= 0.150718
Sum of electronic and zero-point Energies= -906.210957
Sum of electronic and thermal Energies= -906.196589
Sum of electronic and thermal Enthalpies= -906.195644
Sum of electronic and thermal Free Energies= -906.254272

**$^4TS_{2'\text{-}3*}$**
E(scf) = -982.787927272 a.u.
$\nu_{min}$ = -218.4 $cm^{-1}$

| | | | | | | | |
|---|---|---|---|---|---|---|---|
| C | -0.868122 | 0.405606 | -0.064149 | C | -6.975114 | 4.168184 | -0.250821 |
| C | -2.236525 | 0.416613 | -0.150215 | C | -6.293293 | 5.350211 | -0.138861 |
| C | -0.790328 | 2.820529 | 0.127839 | C | -4.855666 | 5.395759 | 0.015036 |
| C | -0.135121 | 1.617787 | 0.076877 | C | -4.157273 | 4.060387 | 0.000882 |
| H | -0.328788 | -0.542404 | -0.103009 | H | -8.058433 | 4.178369 | -0.396350 |
| H | -2.814093 | -0.502798 | -0.256911 | H | -6.823960 | 6.302651 | -0.168010 |
| H | -0.255107 | 3.765449 | 0.233656 | N | -2.841113 | 4.061449 | 0.084937 |
| H | 0.953180 | 1.581774 | 0.143364 | N | -4.294614 | 1.647698 | -0.170139 |
| C | -2.945799 | 1.650582 | -0.097277 | O | -4.212517 | 6.440458 | 0.133628 |
| C | -2.210878 | 2.865189 | 0.040740 | H | -6.569718 | 2.663385 | 1.675360 |
| C | -4.899157 | 2.828782 | -0.118192 | O | -7.133742 | 1.503939 | -1.920727 |
| C | -6.342464 | 2.900979 | -0.133768 | N | -7.039684 | 1.745675 | -0.584314 |

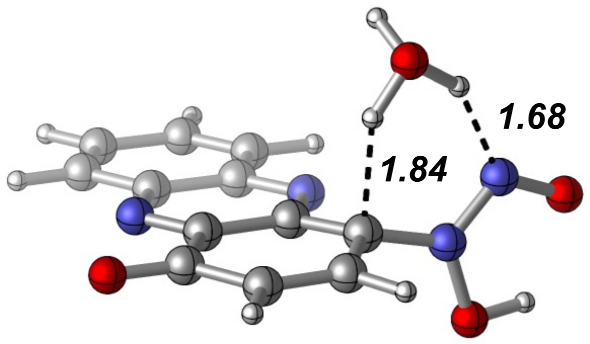


Zero-point correction= 0.217651 (Hartree/Particle)
Thermal correction to Energy= 0.233572
Thermal correction to Enthalpy= 0.234516
Thermal correction to Gibbs Free Energy= 0.173648
Sum of electronic and zero-point Energies= -982.570276
Sum of electronic and thermal Energies= -982.554355
Sum of electronic and thermal Enthalpies= -982.553411
Sum of electronic and thermal Free Energies= -982.61427

**[4]3***

E(scf) = -906.362613200 a.u.

| | | | | | | | |
|---|---|---|---|---|---|---|---|
| C | -3.798412 | -2.117165 | 0.091887 | C | 2.443841 | 1.433233 | -0.097783 |
| C | -2.428547 | -2.158495 | 0.061642 | C | 1.794572 | 2.637446 | -0.051327 |
| C | -3.798324 | 0.307559 | 0.059654 | C | 0.352317 | 2.733126 | -0.013048 |
| C | -4.492363 | -0.873800 | 0.089448 | C | -0.385316 | 1.420392 | -0.025882 |
| H | -4.369939 | -3.046467 | 0.117514 | H | 3.534017 | 1.421968 | -0.175789 |
| H | -1.880214 | -3.101554 | 0.061175 | H | 2.360270 | 3.568918 | -0.081300 |
| H | -4.303651 | 1.274499 | 0.059977 | N | -1.702286 | 1.470486 | 0.013018 |
| H | -5.583003 | -0.869028 | 0.112980 | N | -0.331854 | -1.000159 | -0.014106 |
| C | -1.679726 | -0.947324 | 0.024665 | O | -0.261478 | 3.800455 | 0.016577 |
| C | -2.374964 | 0.297829 | 0.028201 | O | 2.193660 | -1.979765 | -0.946284 |
| C | 0.313957 | 0.158863 | -0.057357 | N | 2.476187 | -0.979080 | -0.106902 |
| C | 1.751225 | 0.200542 | -0.100918 | N | 3.533002 | -1.190313 | 0.611958 |

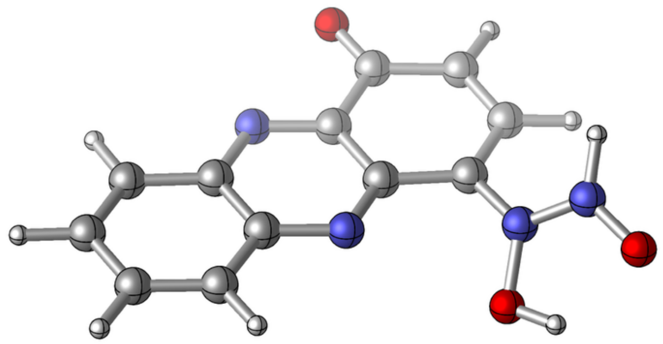

Zero-point correction= 0.194597 (Hartree/Particle)
Thermal correction to Energy= 0.208708
Thermal correction to Enthalpy= 0.209652
Thermal correction to Gibbs Free Energy= 0.152847
Sum of electronic and zero-point Energies= -906.168016
Sum of electronic and thermal Energies= -906.153905
Sum of electronic and thermal Enthalpies= -906.152961
Sum of electronic and thermal Free Energies= -906.209767 9

**$^4TS_{3*-4}$**
E(scf) = -906.351407104 a.u.
$\nu_{min}$ = -580.7 $cm^{-1}$

| | | | | | | | |
|---|---|---|---|---|---|---|---|
| C | 0.707285 | -0.548052 | -0.031334 | C | -5.509443 | 3.050610 | 0.344015 |
| C | -0.655741 | -0.577919 | 0.125998 | C | -4.869757 | 4.235831 | 0.157769 |
| C | 0.710261 | 1.869719 | -0.234639 | C | -3.422704 | 4.325172 | -0.001592 |
| C | 1.396552 | 0.682197 | -0.212197 | C | -2.675613 | 3.019796 | 0.049722 |
| H | 1.274591 | -1.479692 | -0.017473 | H | -6.591215 | 3.039284 | 0.491749 |
| H | -1.198644 | -1.513169 | 0.266599 | H | -5.424809 | 5.174165 | 0.143515 |
| H | 1.214179 | 2.826885 | -0.372411 | N | -1.370758 | 3.052481 | -0.103947 |
| H | 2.480297 | 0.674633 | -0.333909 | N | -2.730440 | 0.625384 | 0.267082 |
| C | -1.386981 | 0.637851 | 0.107316 | O | -2.831205 | 5.385229 | -0.160649 |
| C | -0.701615 | 1.874325 | -0.075939 | O | -4.981000 | -0.493069 | 0.731163 |
| C | -3.378361 | 1.783433 | 0.243695 | N | -5.568630 | 0.684927 | 0.537262 |
| C | -4.820652 | 1.795864 | 0.389687 | N | -6.800768 | 0.370344 | -0.672114 |

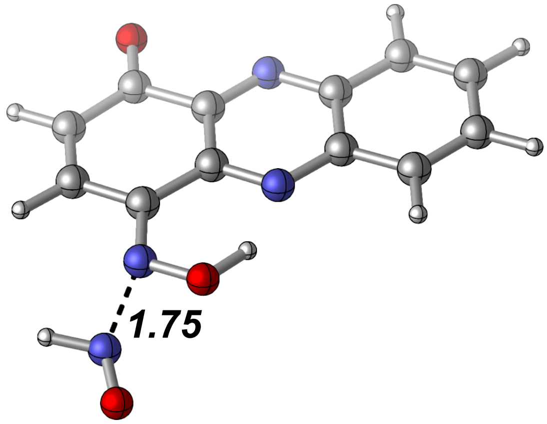


Zero-point correction= 0.190551 (Hartree/Particle)
Thermal correction to Energy= 0.204780
Thermal correction to Enthalpy= 0.205724
Thermal correction to Gibbs Free Energy= 0.148488
Sum of electronic and zero-point Energies= -906.160856
Sum of electronic and thermal Energies= -906.146627
Sum of electronic and thermal Enthalpies= -906.145683
Sum of electronic and thermal Free Energies= -906.20
